# Influence of genetic variants on gene expression in human pancreatic islets – implications for type 2 diabetes

**DOI:** 10.1101/655670

**Authors:** Ana Viñuela, Arushi Varshney, Martijn van de Bunt, Rashmi B. Prasad, Olof Asplund, Amanda Bennett, Michael Boehnke, Andrew Brown, Michael R. Erdos, João Fadista, Ola Hansson, Gad Hatem, Cédric Howald, Apoorva K. Iyengar, Paul Johnson, Ulrika Krus, Patrick E. MacDonald, Anubha Mahajan, Jocelyn E. Manning Fox, Narisu Narisu, Vibe Nylander, Peter Orchard, Nikolay Oskolkov, Nikolaos I. Panousis, Anthony Payne, Michael L. Stitzel, Swarooparani Vadlamudi, Ryan Welch, Francis S. Collins, Karen L. Mohlke, Anna L. Gloyn, Laura J. Scott, Emmanouil T. Dermitzakis, Leif Groop, Stephen C.J. Parker, Mark I. McCarthy

## Abstract

Most signals detected by genome-wide association studies map to non-coding sequence and their tissue-specific effects influence transcriptional regulation. However, many key tissues and cell-types required for appropriate functional inference are absent from large-scale resources such as ENCODE and GTEx. We explored the relationship between genetic variants influencing predisposition to type 2 diabetes (T2D) and related glycemic traits, and human pancreatic islet transcription using RNA-Seq and genotyping data from 420 islet donors. We find: (a) eQTLs have a variable replication rate across the 44 GTEx tissues (<73%), indicating that our study captured islet-specific *cis*-eQTL signals; (b) islet eQTL signals show marked overlap with islet epigenome annotation, though eQTL effect size is reduced in the stretch enhancers most strongly implicated in GWAS signal location; (c) selective enrichment of islet eQTL overlap with the subset of T2D variants implicated in islet dysfunction; and (d) colocalization between islet eQTLs and variants influencing T2D or related glycemic traits, delivering candidate effector transcripts at 23 loci, including *DGKB* and *TCF7L2*. Our findings illustrate the advantages of performing functional and regulatory studies in tissues of greatest disease-relevance while expanding our mechanistic insights into complex traits association loci activity with an expanded list of putative transcripts implicated in T2D development.

## Introduction

Genome-wide association studies (GWAS) have generated a growing inventory of genomic regions influencing type 2 diabetes (T2D) predisposition and related glycemic traits ^1,2^. However, progress in defining the mechanisms whereby these associated variants mediate their impact on disease-risk has been slow^3^. Over 90% of the associated signals map to non-coding sequence^4,5^ complicating efforts to connect T2D-associated variants with the transcripts and networks through which they exert their effects. One approach for addressing this “variant-to-function” challenge is to use expression quantitative trait loci (eQTL) mapping to characterize the impact of disease-associated regulatory variants on the expression of nearby genes^6^.

Demonstrating that a disease-risk variant co-localizes with a cis-eQTL signal is consistent with a causal role for the transcript concerned, a hypothesis that can then be subject to more direct evaluation, for example, by perturbing the gene in suitable cellular or animal models. However, eQTL signals are often tissue-specific^7^: consequently, the power to detect mechanistically-informative expression effects is dependent on assaying expression data from sufficient numbers of samples across the range of disease-relevant tissues^6^.

The pathogenesis of T2D involves dysfunction across multiple tissues, most obviously pancreatic islets, adipose, muscle and liver. Risk variants that influence T2D predisposition through processes active in each of these have been reported (e.g. *KLF14* in adipose^8^, *TBC1D4* in muscle^9^, *ADCY5* in islets^10^, *GCKR* in liver^11^). However, multiple physiological and genomic analyses consistently indicate that islet dysfunction makes the greatest contribution to T2D risk ^4,12,13^. Research access to human pancreatic islet material is therefore essential, and previous studies have demonstrated the potential of islet expression information to characterize T2D effector genes such as *MTNR1B* and *ADCY5*^14-16^. However, access to human islet material is limited, and the largest published human islet RNA-Seq dataset includes only 118 samples^16^.

We constituted the InsPIRE (**I**ntegrated **N**etwork for **S**ystematic analysis of **P**ancreatic **I**slet **R**NA **E**xpression) consortium as a vehicle for the aggregation and joint analysis of human islet RNA-Seq data^15-17^. Here, we report analyses of 420 human islet preparations which provide a detailed landscape of the genetic regulation of gene expression in this key tissue, and its relationship to mechanisms of T2D predisposition.

Our research addresses questions with relevance beyond T2D. When a disease-relevant tissue is missing from reference datasets such as GTEx, what additional value accrues from dedicated expression profiling from that missing tissue? What is the impact of tissue heterogeneity on the interpretation of eQTL data? What does the synthesis of tissue specific epigenomic and expression data tell us about the coordination of upstream transcription factor regulators of gene expression? And, finally, what information do tissue-specific eQTL analyses provide about the regulatory mechanisms mediating disease predisposition?

### Characterization of genetic regulation of gene expression in islets

We combined islet RNA-Seq with dense genome-wide genotype data from 420 individuals. Data from 196 of these individuals have been reported previously^14-17^. We aggregated, and then jointly mapped and reprocessed, all samples (median sequence-depth per sample ∼60M reads) to generate exon- and gene-level quantifications, using principal component methods to correct for technical and batch variation (Methods; Supp. Figure SF1).

To characterize the regulation of gene expression for the 17,914 protein coding and long non-coding RNAs (lncRNAs) genes with quantifiable expression in these samples, we performed eQTL analysis (fastQTL^18^) on both exon and gene-level expression measures, using all 8.05M variants that pass quality control (QC) (Methods; Supp. Tables ST1 & ST2). This joint analysis of all 420 individuals identified 4,312 genes (eGenes) with significant *cis-*eQTLs at the gene level (FDR<1%; *cis* defined as within 1Mb of the transcription start site [TSS]). The complementary exon-level analysis, which can capture the impact of variants influencing splicing as well as expression, detected 6,039 eGenes (FDR<1%, Supp. Figure SF2)^19,20^. Stepwise regression analysis (after conditioning on the lead variant) identified a further 1,702 independent eQTLs (involving 1,291 eGenes), giving a total of 7,741 islet exon-level eQTLs (Supp. Tables ST1 & ST2). At the 1,291 eGenes with at least two independent exon-eSNPs, although primary eSNPs tended to localize closer to the canonical TSS than secondary eSNPs (Wilcoxon test *P*=6.3×10^-30^), there were 503 (39.0%) of these genes for which the second eSNP identified during stepwise conditional analysis was more proximal to the TSS (Supp. Figure SF2).

### Tissue specific regulatory variation in islets

For many complex traits of biomedical interest, the value of targeting the specific cell-types of interest for dedicated eQTL discovery -- as opposed to relying on existing eQTL data from more accessible tissues – remains unclear. To examine this, we considered the degree to which the set of 6,039 exon-level islet eQTLs overlapped eQTLs detected in 44 tissues (N>70) from version 6p of GTEx^7^. To allow direct comparison with InsPIRE, we reprocessed GTEx data to generate exon-level eQTLs (Methods). Of the 6,039 islet eGenes, 5% (337) had no significant eQTLs (including both exon- or gene-level analyses from GTEx) in any of the 44 tissues (Supp. Table ST3). Instead of defining “tissue-sharing” using arbitrary thresholds, we used P-value enrichment analysis (π_1_) ^21^ to measure the proportion of islet eQTLs shared with other GTEx tissues: estimates ranged from 40% (hypothalamus) to 73% (adipose). We saw the expected positive linear relationship between π_1_ measures and sample sizes for the respective tissues in GTEx^7^ (**Figure 1A**). However, π_1_ reached only 65% and 57% (respectively) for skeletal muscle (n=361), and whole blood (n=338), the tissues with the largest representation in this version of GTEx. On the other hand, whole pancreas, often naively-used as a surrogate for the T2D-relevant islet component, represents an imperfect proxy for islet (π_1_=0.67 with islets). This does not reflect low sample size: the number of whole pancreas samples is on a par with other tissues such as skin and spleen with comparable eQTL-sharing (π_1_ 0.67, 0.61 respectively). These data demonstrate that there is a component of tissue-specific genetic regulation that could, at these sample sizes, only be detected in islets, illustrating the value of extending current expression profiling efforts to additional tissues and cell-types of particular biomedical importance. This also indicates that whole pancreas has no particular advantage as a proxy for islet eQTLs.

**Figure 1.**
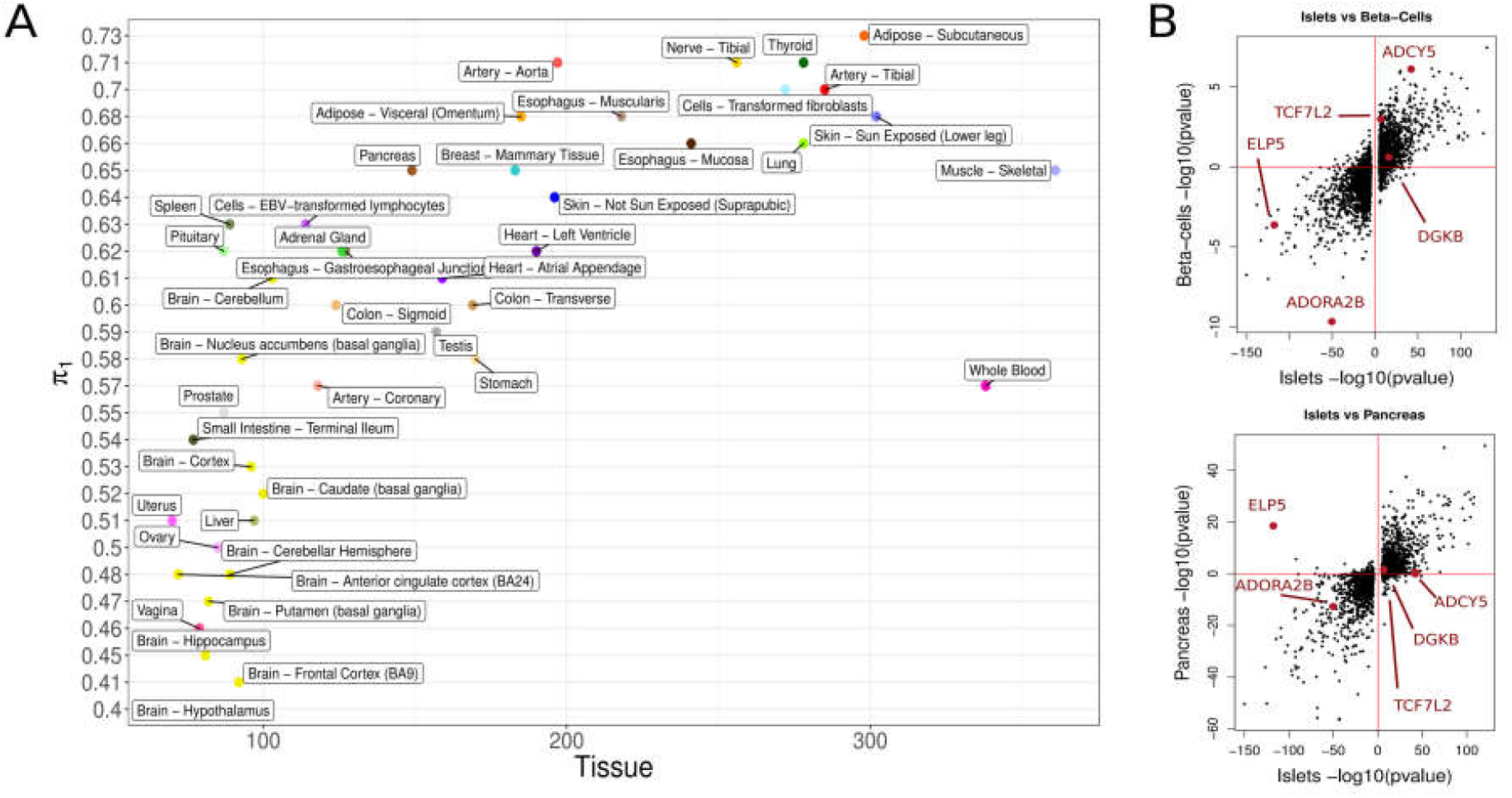
Islet eQTL discovery. **A)** Proportion of islet eQTLs active in GTEx tissues using *P*-value enrichment analysis (π_1_ estimate for replication). **B)** Comparison between eQTLs discovered in islets and their pvalues in beta-cells (top figure, N=26) and whole pancreas tissue from GTEX (bottom figure, n=149). The axes show the - log10 Pvalue of the eQTL associations adjusted by the eQTL direction of effect with respect to the reference allele.

### Cellular heterogeneity

The human islets analyzed in this, and other, studies include a mixture of cell types, including the hormone-producing α, β and δ-cells, and a variable amount of adherent exocrine material. From the perspective of T2D pathogenesis, the transcriptomes of the former are of most interest. However, the eQTLs identified could have their origins from any of the cellular components. We used a number of approaches to address interpretative challenges resulting from this cellular heterogeneity.

First, we performed tissue deconvolution analysis to estimate the proportion of exocrine contamination: these analyses were performed prior to the principal component adjustment used to generate the main results and used reference expression signatures for exocrine pancreas, beta-cells and islet non-beta-cells, the last two from a subset (n=26) of the islet preparations FAC-sorted using the zinc-binding dye Newport Green^17^ (Methods). Estimates of the proportion of exocrine pancreas contamination ranged from 1.8% to 91.8% (median 33.5%): these were significantly correlated (r=0.50, *P*=2.8×10^-15^) with independent estimates of exocrine content obtained at islet collection by dithizone staining (n=232) (Supp. Figure SF3). Within the islet endocrine fraction, median estimates of beta-cell (58.8%, IQR 43.2-66.9%) and non-beta-cell (41.2%, 33.1-56.8%) fractions are in agreement with estimates from morphometric assessment ^22^. In 34 samples from donors annotated as having T2D, median estimates of beta-cell composition were lower than those from non-diabetic donors (n=330) (31.8% vs. 35.6%, *P*=4.5×10^-4^, Supp. Figure SF3). This provides independent confirmation, based on transcriptomic signatures, of evidence, from morphometric and physiological studies, that the functional mass of beta-cells is reduced in T2D^23,24^.

Of the 420 InsPIRE samples, beta-cell enriched transcriptomes were available for 26 following FAC-sorting. With this limited sample size, the only eQTL reaching significance and then only at a less stringent threshold of FDR<5% (Supp. Table ST4) was at *ADORA2B* (*P*=3.8×10^-10^, beta=-1.20): this signal was also detected in InsPIRE islets (*P*=3.9×10^-51^, beta=-0.65) and GTEx pancreas (*P*=1.6×10^-16^, beta=-0.73) (Supp. Figure SF4; Supp. Table ST5). By comparing the p-value distributions of the eQTLs in islets vs beta-cells^21^, we estimate that 46% of islet eQTLs are active in beta-cells (Figure 1B). By extracting beta-cell association results from the 7,741 independent SNP-exon pairs significant in islets, 227 were also significant in beta-cells (FDR<1%, Supp. Table ST6). Genes with cell-type-specific regulatory effects were sought by testing for interactions between genotype and cellular fraction estimates, controlling for technical variables (Methods). We identified 18 islet *cis*-eQTLs with a “genotype-by-beta-cell proportion” interaction and 8 with a “genotype-by-exocrine cell proportion” interaction (FDR<1%, Supp. Tables ST7, ST8 & ST9).

We conclude that a substantial proportion of the regulation of gene expression detected in pancreatic islets derives from cell-specific effects. Ongoing efforts to develop a single-cell view of islet transcriptional signatures should inform these analyses, although the limited sample size of current studies^25-28^ and the lack of genotype information means they offer little direct insight into the relationship between genetic variation and cell-type-specific transcript abundance.

### Functional properties of islet genetic regulatory signals

Using previously-published islet chromatin states derived from histone modification data^14^, we observed a significant enrichment of islet eSNPs in active islet chromatin states including active TSS (fold enrichment=3.84, *P*=5.5×10^-206^), active enhancers (fold enrichment>1.73, *P*<4.8×10^-04^ between two enhancer states) and stretch enhancers (fold enrichment=1.57, *P*=2.7×10^-13^), with concomitant depletion of eSNPs in repressed and quiescent states (fold enrichment <0.66) (Supp. Table ST10; Supp. Figure SF5). This recapitulates the enrichment observed for T2D GWAS signals within active islet chromatin (Supp. Figure SF6)^10,14,29,30^. Next, we examined the relationship between the chromatin context of islet eSNPs and their effect sizes (**Figure 2A**): eSNPs that overlap active TSS chromatin states had larger effects than those in repressed or weak-repressed polycomb states (Wilcoxon Rank Sum Test *P*=0.039).

**Figure 2.**
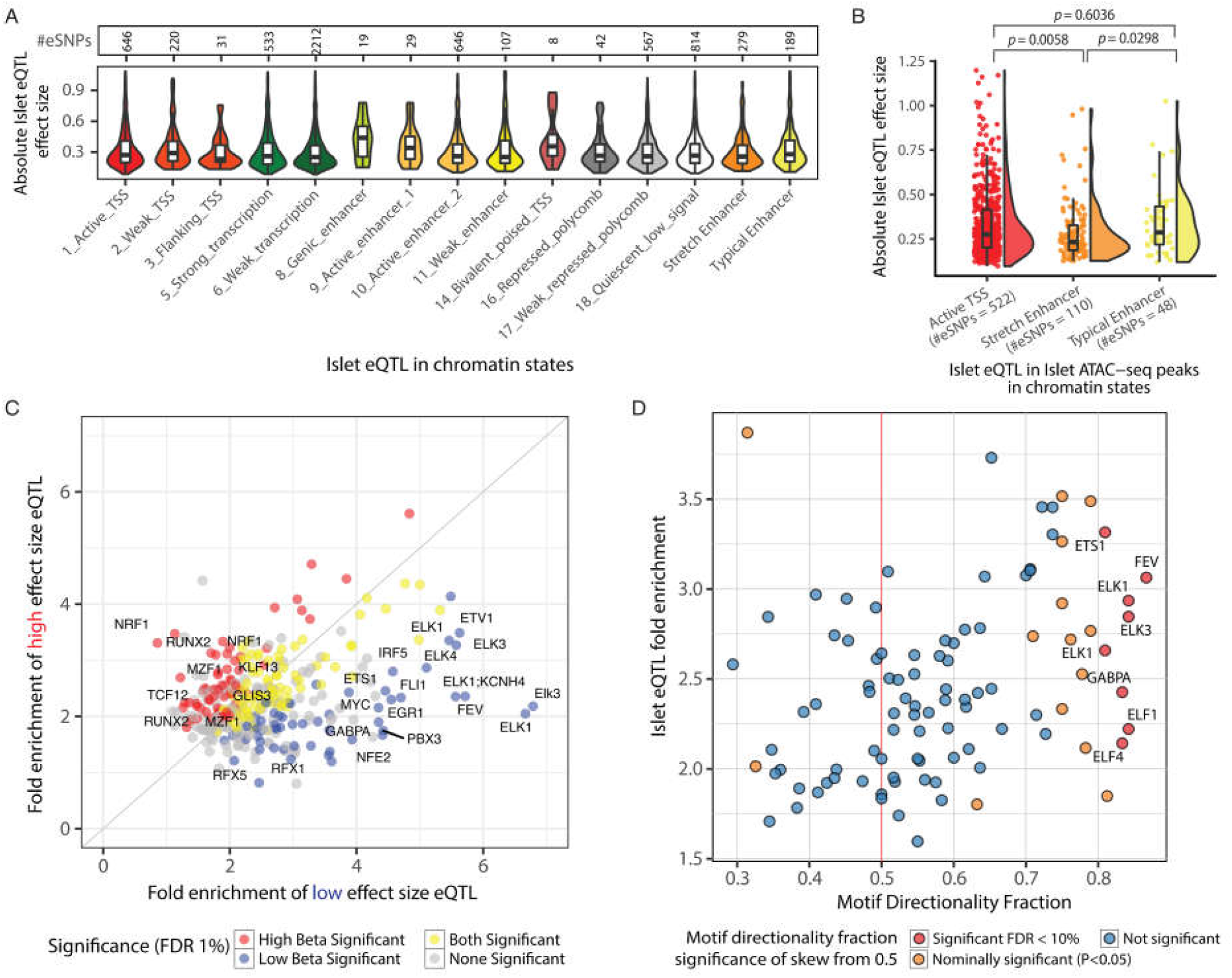
Integration of Islet eQTL with epigenomic information reveals characteristics of gene expression regulation. **A)** Distribution of absolute effect sizes for Islet eQTLs in each Islet chromatin state. **B)** Distribution of absolute effect sizes for Islet eQTL in ATAC-seq peaks in three Islet chromatin states. eQTL SNPs in ATAC-seq peaks in stretch enhancers have significantly lower effect sizes than SNPs in ATAC-seq peaks in active TSS and typical enhancer states. P values obtained from a Wilcoxon rank sum test. **C)** Fold Enrichment for transcription factor footprint motifs to overlap low vs high effect size islet eQTL SNPs. **D)** TF footprint motif directionality fraction vs fold enrichment for the TF footprint motif to overlap islet eQTL. TF footprint motif directionality fraction is calculated as the fraction of eQTL SNPs overlapping a TF footprint motif where the base preferred in the motif is associated with increased expression of the eQTL eGene. Significance of skew of this fraction from a null expectation of 0.5 was calculated using Binomial test.

Because chromatin states represent integrated histone mark patterns, and transcription factors (TFs) are more likely to bind open DNA, we next considered regions of accessible chromatin within each of the chromatin states, using human islet ATAC-seq data^14^. As expected, a high proportion (80%) of islet eQTLs (based on the lead eSNP or proxies [r^2^>0.99]) overlap islet ATAC-seq peaks in islet-active TSS chromatin states: 511 (80.8%) of 646 islet eSNPs overlapping islet active TSS chromatin lie in the (ATAC-defined) open chromatin portion of that chromatin state (Supp. Figure SF7). Almost half (49.7%) the islet-active TSS chromatin state territory is occupied by islet ATAC-seq peaks. When we considered only those eSNPs within islet ATAC-seq peaks, those within stretch enhancers (islet-specific enhancer chromatin state segments >3kb^29^) had smaller effects than those in either typical enhancers or active TSSs (Wilcoxon Rank Sum Test *P*=0.0088, *P*=0.0099, respectively) (**Figure 2B**).

One corollary is that eSNPs in cell-specific stretch enhancers are, for equivalent effect size, likely to require larger sample size studies for eQTL discovery than those in other annotations.

We previously reported enrichment of selected TF footprint motifs at islet eSNPs^14^. Here, with a larger eSNP catalog, we sought to determine how eSNP effect size and target gene expression directionality is associated with base-specific TF-binding preferences. Using published TF footprint data (*in vivo*-predicted TF motif binding) from human islet ATAC-seq analyses^14^, we partitioned eSNPs into two equally-sized bins (absolute slope ≥ vs. <0.254 standard deviation units). Higher effect-size eSNPs were preferentially enriched (<1% FDR) for footprint motifs characteristic of islet-relevant TF families, including KLF11 (motif=KLF13_1, *P*=5.3×10^-6^) and GLIS3 (motif GLIS3_1, P=5.2×10^-6^). Other footprint motifs, including the RFX and ETS families of TFs, were significantly enriched for low effect-size eSNPs (*P*<2×10^-4^) (**Figure 2C**, Supp. Table ST11).

Finally, since TFs can act as activators, repressors, or both^31^, we asked, using previously-published MPRA (massively parallel reporter assay) data from HepG2 and K562 cell lines^32^., whether eSNP alleles matching the base preference at TF footprint motifs have a consistent directional impact on gene expression. We defined a motif directionality fraction score (ranging from repressive [0] to activating [1]) for each TF footprint motif (Methods). Of the 99 motifs reported as consistently activating or repressive across HepG2 and K562 cell lines that were present in our study, only 8% (n=8) showed skewed activator preference in islets (<10% FDR; Figure 2D, Supp. Table ST12). The activator motifs we identified include many ETS family members which have a known preference for transcriptional activation^32^.

Our analyses demonstrate the value of contrasting tissue-specific stretch enhancers with more ubiquitous TSS states to delineate the role of underlying chromatin on function; and illustrate how the integration of eQTL information with ATAC-seq and high-resolution TF footprinting reveals the *in vivo* activities of these upstream regulators.

### Islet eQTLs are enriched among T2D and glycemic GWAS variants

Diverse lines of evidence emphasize the contribution of reduced pancreatic islet function to the development of T2D, with many T2D GWAS loci act primarily through reducing insulin secretion ^4,10,12,33^. To examine the relationships between tissue-specific regulation of gene expression and T2D predisposition alleles, we focused on 78 lead GWAS SNPs with the strongest associations to T2D (as reported in Fuchsberger et al.^3^) and 44 variants significantly associated with T2D-relevant continuous glycemic traits, including fasting glucose and beta-cell function (HOMA-B) in non-diabetic individuals (Supp. Table ST13)^34-36^. For comparison, we included 55 GWAS variants implicated in T1D-predisposition^37^. To determine the extent to which the GWAS variants were selectively enriched for islet eQTL associations, we extracted exon-level eQTL information for each of these variants from InsPIRE and the 44 GTEx^7^ tissues. We compared observed effect-size estimates to those derived from a null distribution of 15,000 random eSNPs, matched to the GWAS SNPs with respect to the number of SNPs in LD, distance to TSS, number of nearby genes and minor allele frequency (Methods). Figure 3A shows the enrichment in eQTL effect sizes at T2D/glycemic GWAS-associated variants for five tissues implicated in T2D pathogenesis (subcutaneous adipose tissue, skeletal muscle, liver, islets, plus hypothalamus), with pancreas and whole blood for comparison.

**Figure 3.**
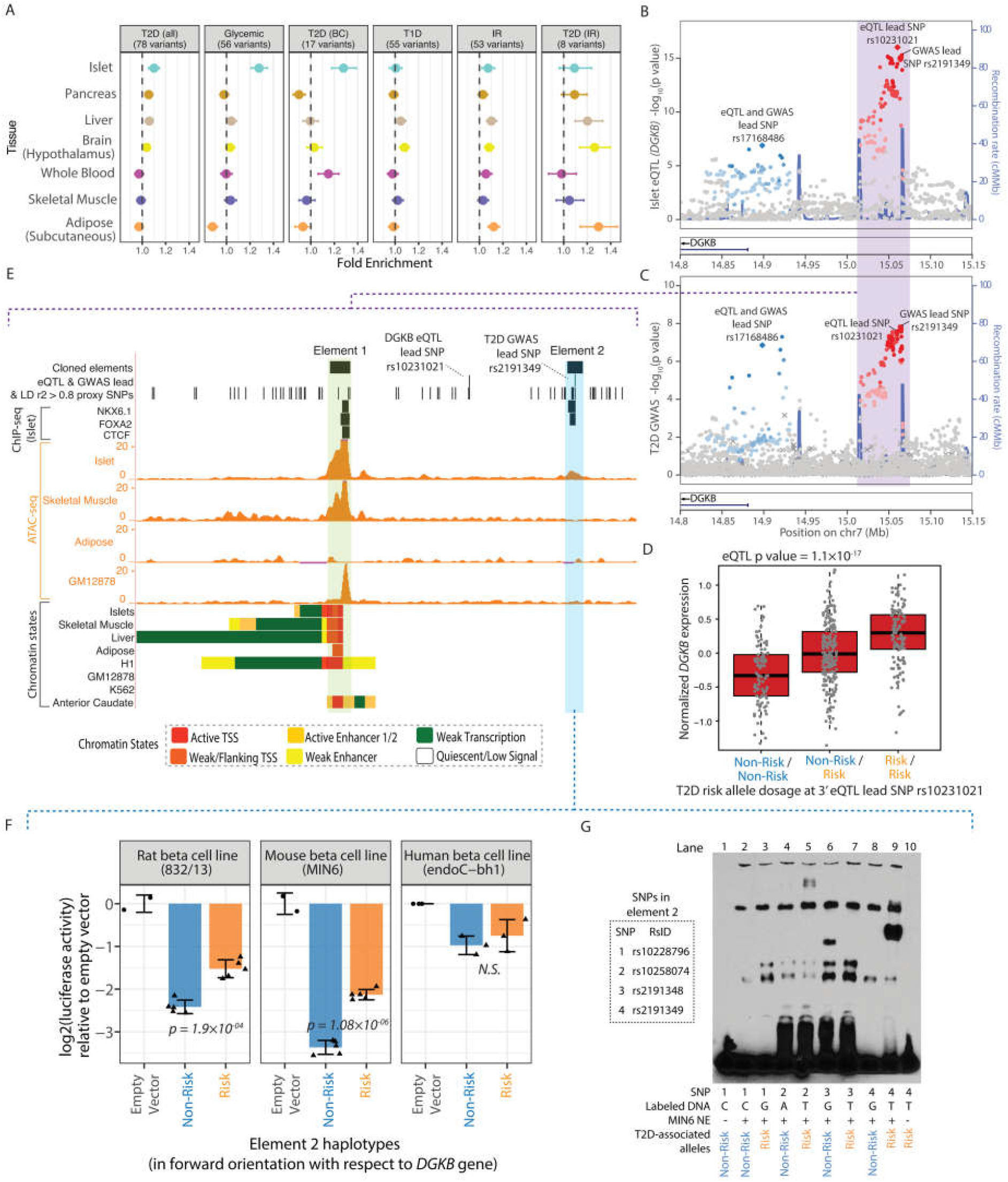
Functional validation of *DGKB* eQTL locus. **A)** Enrichment of eQTL effect sizes in different GTEx tissues at T2D/glycemic GWAS-associated variants. Numbers within square brackets denote the number of variants implicated for the trait. Also shown are subsets of T2D GWAS associated with reduced insulin secretion or islet beta cell dysfunction (T2D (BC)) or insulin resistance (T2D (IR)), type 1 diabetes (T1D) signals, insulin resistance (IR). **B)** Two independent islet eQTL signals (lead SNP rs17168486 referred at as the 5’ signal and lead SNP rs10231021 referred to as the 3’ signal) are identified near the *DGKB* gene locus. These signals co-localize with two independent T2D GWAS signals shown in **C)** where (rs17168486 referred to as the 5’ signal and lead SNP rs2191349 referred to as the 3’ signal and. LD information was not available for SNPs denoted by (×). **D)** Normalized *DGKB* gene expression levels relative to the T2D risk allele dosage at the 3’ islet eQTL for *DGKB* lead SNP rs10231021. eQTL P value adjusted to the beta distribution is shown. **E)** Genome browser view of the region highlighted in purple in (B) and (C) that contains the 3’ *DGKB* eQTL and T2D GWAS signals. Two regulatory elements overlapping islet ATAC-seq peaks (element 1 highlighted in green, element 2 highlighted in blue) were cloned into a luciferase reporter assay construct for functional validation. **F) L**og 2 Luciferase assay activities (normalized to empty vector) in rat (832/13), mouse (MIN6) and human (endoC) beta cell lines for the element 2 highlighted in blue in (**D**). Risk haplotype shows significantly higher (P<0.05) activity than the non-risk haplotype in 832/13 and MIN6, consistent with the eQTL direction shown in (**F**). P values were determined using unpaired two-sided t-tests. **G)** Electrophoretic mobility shift assay (EMSA) for probes with risk and non-risk alleles at the four SNPs overlapping the regulatory element validated in (**F**) using nuclear extract from MIN6 cells.

We detected significant enrichment for islet eQTLs for variants associated with continuous glycemic traits (normalize enrichment score (NES)=1.27; *P*=3.6×10^-3^) (Supp. Figure SF11, Supp. Table ST14): apart from a modest signal in ovary (NES=1.13, *P*=0.02), there was no enrichment in other GTEx tissues. Islet eQTL enrichment for the 78 T2D variants was directionally consistent but did not reach nominal significance (NES=1.10; *P*=0.07). However, T2D GWAS signals act through physiological effects in multiple tissues^8,9^, and significant enrichment for islet eQTL signals (NES=1.27; p=0.025) was seen in the subset (n=17) of T2D GWAS signals for which the evidence (based on the patterns of association to other T2D-related traits) points most clearly to mediation through reduced insulin secretion (beta-cell dysfunction (BC))^4,10,12,33^ For this subset, there was no enrichment for eQTL effect sizes in whole pancreas (NES=0.90, *P*=0.88). No enrichment of islet eQTL signals was seen for the T1D-risk variants, consistent with the consensus that genetic risk for T1D is largely mediated through immune mechanisms^37^. In the subset of eight T2D GWAS signals with the strongest evidence of mediation through insulin resistance (IR), eQTL enrichment was detected in insulin-target tissues such as liver (NES=1.10; *P*=0.03) and adipose tissue (NES=1.12; *P*=0.04), but not in islets (NES=1.07, *P*=0.17). Similar patterns of eQTL enrichment were seen for a broader, partly-overlapping, set of 53 lead variants influencing insulin sensitivity derived from a multivariate GWAS^38^.

These data reveal tissue-specific patterns of genetic regulatory impact for variants at T2D- and glycemic-trait loci which mirror the mechanistic inferences generated by physiological analysis of those signals. They also highlight the importance of matching the tissue origin of the transcriptomic data used for mechanistic inference to the tissue-specific impact of each GWAS signal on disease predisposition.

### Identifying effector transcripts for T2D and glycemic traits

Previous studies have identified GWAS signals displaying apparent overlap between islet eQTLs and the T2D/glycemic GWAS signals^14-16^, but not all of these signals have been evaluated with respect to the statistical evidence for co-localization and not all coincident signals have replicated despite ostensibly similar designs and power^16^.

There are multiple methods for evaluating the evidence for co-localization: these make different assumptions and often lead to discrepant results^39^. We focused on the co-localization evidence provided by two complementary algorithms: COLOC^40^, which assesses differences in regression coefficients of variants associated to two traits, and RTC^41^, which assesses the differences in ranking of SNPs associated with one trait after conditioning on the most associated SNP for the other. We detected evidence for co-localization (using either method) for islet eQTLs at 23 GWAS loci, (comprising 24 independent signals, given two signals at *DGKB*) 16 of them reflecting T2D associations (Supp. Table ST15).

Evidence for co-localization was most compelling for 11 loci (12 signals) at which both RTC and COLOC provided strong support: including extending confirmation of previously observed co-localizations at *ADCY5, HMG20A, IGF2BP2* and *DGKB*^10,42^ as candidate effector transcripts. At other loci, we observed islet *cis-*eQTL co-localization for the first time. For example, previous efforts to characterize the mechanism of action at the *TCF7L2* locus have demonstrated that the fine-mapped T2D-risk allele at rs7903146 influences chromatin accessibility and enhancer activity in islets^43^, but evidence linking these events to *TCF7L2* expression has been missing. Our data reveals that the rs7309146 increases islet expression of *TCF7L2* (eQTL beta=0.21, *P*=1.9×10^-7^) (Supp. Figure SF12). The same eQTL signal was also detected in the smaller beta-cell specific analysis (n=26; eQTL beta= 0.72; *P*=1.0×10^-3^). The association between rs7903146 and *TCF7L2* expression was restricted to islets, consistent with evidence that non-diabetic carriers of the *TCF7L2* risk-allele display markedly reduced insulin secretion^44^. Recent studies have proposed *ACSL5*^45^ as a possible effector transcript at this locus, but we found no support - in any tissue - that rs7903146 influences *ACSL5* expression.

Several expected signals of GWAS/islet-eQTL overlap were not observed in our exon-eQTL based analysis. *MTNR1B* has shown consistent islet cis-eQTL signals in previous studies^16,46^, but was excluded from our exon-level analysis due to low exonic-read coverage: in the complementary gene-level analyses, there was strong evidence of co-localization between the lead T2D variant (rs10830963) and *MTNR1B* expression (*P*=5.3×10^-21^; Supp. Table ST2). At *ZMIZ1*, the previously-reported *cis-*eQTL was nominally significant (rs185040218; *P*=3.0×10^-5^) but did not reach the 1% FDR threshold for inclusion in co-localization testing.

At other loci, complex, but divergent, patterns of association between the eQTL and T2D GWAS signals challenged the assumptions of these co-localization methods. At the *ZBED3* locus, for example, the association plots highlight two distinct T2D signals (∼500kb apart), and two distinct islet-eQTL signals for *PDE8B*, but only the signal at rs7708285 appears coincident (Supp. Figure SF13). COLOC detects this as co-localization, but this configuration cannot easily be tested using RTC which restricts analysis to a single haplotype block.

Finally, we attempted to characterize eGenes that overlapped T2D/glycemic GWAS signals by assessing the impact of changes in glycemic status on islet expression. We used data from a recent analysis of islets recovered from diabetic and non-diabetic donors, focussing on transcripts that showed acute changes in expression when exposed to glucose levels in culture that contrasted with those to which they had been habituated^47^ (Supp. Table ST16). Islet eGenes such as *STARD10, WARS, SIX3, NKX6-3* and *KLHL42* which may be of particular interest in that their expression in islets is regulated both by T2D-associated variation and by acute changes in glucose exposure.

### Experimental validation at *DGKB*

The *DGKB* locus features two independent GWAS signals and three independent eQTLs. Only two of these show islet eQTL co-localizations: at both, the T2D-risk allele is associated with increased islet expression of *DGKB* (Figure 3B, 3C and 3D). Physiological analyses for these variants are consistent with mediation through islet dysfunction^4,33^.

At the 3’ signal, the lead eSNP, rs10231021 (Figure 3B), is in high LD (r^2^=1, D’=1) with the lead GWAS variant, rs2191349 (Figure 3C). For functional follow-up, we considered seven variants that mapped to islet ATAC-Seq peaks and were in high LD (r^2^>0.8) with rs2191349 (Figure 3E). Three (rs7798124, rs7798360, rs7781710, Figure 3D, “Element 1”) overlap an ATAC-seq peak shared across islets, skeletal muscle and the lymphoblastoid cell line GM12878^48^ cell-line: four others (rs10228796, rs10258074, rs2191348, rs2191349, Figure 3E “Element 2”) lie in a smaller but more islet-specific peak. We cloned these putative regulatory elements into luciferase reporter constructs and performed transcriptional reporter assays in three widely-used cellular models of the beta-cell (human EndoC-βH1^49^, rat INS1-derived 823/13 (Methods), mouse MIN6^50^. Element 1 showed consistent enhancer activity across all three lines but no allelic differences consistent with the eQTL direction of effect (Supp. Figure SF14). Element 2 showed reduced luciferase expression in all three beta-cell lines when in forward orientation with respect to *DGKB*. The T2D-risk haplotype showed higher expression than non-risk in 832/13 (P=1.9×10^-4^) and MIN6 (P=1.1×10^-6^). Equivalent experiments in EndoC-βH1 showed a consistent trend, which did not reach significance (Figure 3F). Luciferase assays using element 2 in reverse orientation also showed consistent trends across the cell lines, reaching significance in 832/13 (Supp. Figure SF14). In electrophoretic mobility assays using MIN6 nuclear extract, three “Element 2” variants (rs10228796, rs2191348, rs2191349) showed allele-specific binding (Figure 3G), supporting a functional regulatory role. These data suggest that T2D-risk alleles alleviate regulatory element repression and are directionally-consistent with the 3’ *DGKB* eQTL (Figure 3D).

At the 5’ eQTL, we focused attention on rs17168486, which was both the lead SNP for islet *cis-* expression and T2D-association, and located in an islet ATAC-seq peak (Supp. Figure SF14). However, luciferase reporter constructs found no consistent allelic effects on transcriptional activity (Supp. Figure SF14).

## Discussion

We have used transcriptome sequencing in 420 human islet preparations to address issues of general relevance to the mechanistic interpretation of non-coding association signals detected by GWAS. We documented the degree to which RNA-sequencing of a disease-relevant tissue missing from a reference set (e.g. GTEx) provides a more complete survey of islet eQTLs. We used this information to extend the number of association signals for T2D and related glycemic traits co-localizing with islet eQTLs, identifying novel candidate effector transcripts at several loci. We explored how cellular heterogeneity (both within the tissue of interest, and reflecting contamination with cells not of direct relevance) can complicate the interpretation of GWAS signal colocalization. We integrated our eQTL catalogue with islet epigenomic data to reveal effect size heterogeneity attributable to local chromatin context and to infer *in vivo* TF directional activities.

Analyses of the physiological association patterns and regulatory annotation enrichment signals of T2D-risk alleles indicate that many, though by no means all, act through the islet^8,9,11,13,51^. A major motivation behind development of this enhanced catalog of islet eQTLs was to support identification of effector transcripts mediating the downstream consequences of these non-coding alleles. At *DGKB*, for example, evidence that both the T2D signals co-localize with islet eQTLs with directionally-consistent impacts on *DGKB* expression lends credibility to a causal role for *DGKB* in T2D-predisposition.

However, it is important to emphasize that robust inference from the coincidence of eQTLs and GWAS signals is difficult. First, the expression data in our study are derived from human islets cultured in basal glycemic conditions: eQTL signals restricted to a subset of the cells within those islets would have been hard to detect, and the same for genes whose expression is dependent on stimulation. Since not all T2D loci act through the mature islet, some of the eQTLs detected may reflect tissue-specific regulation that is not germane to the development of the diabetic phenotype. At some loci, this may reflect variants that influence T2D-risk through effects on islet development. Reassuringly, for the co-localizing loci we detected, our analyses – including selective enrichment of islet eQTLs in the subset of T2D loci that primarily influence insulin secretion - are consistent with mediation through islet dysfunction.

Second, confident assignment of co-localization can be difficult. There are multiple algorithms to assess the evidence that two association signals are likely to reflect the same causal variants, but agreement between them is incomplete^39^. An additional challenge arises from the complex architecture of many GWAS signals, such that conditional decomposition is required before co-localization across multiple overlapping signals can be accurately assigned^52^. This is especially important when the sets of GWAS and *cis-*eQTL signals at a given locus are not completely overlapping, since obvious co-localization at one of the contributing signals can be masked by differences in the overall shape of the association signals that confounds simplistic analysis.

Third, recent studies have shown that functionally-constrained genes – which are depleted for missense or loss-of-function variants – are also less likely to have eQTLs, indicating uniform intolerance of both regulatory and coding variation^53,54^. Complementary studies focusing on regulatory elements have shown that large, cell-specific stretch enhancers harbor smaller effect size eQTLs than ubiquitous promoter regions^55^ and that genes with more cognate enhancer sequence are depleted for eQTLs^56^. Our findings that islet eQTLs that map to the islet stretch enhancers most frequently implicated in GWAS regions have smaller eQTL effect sizes is consistent with these observations. One consequence is that, when a GWAS variant has regulatory impact on multiple *cis*-genes, eQTL-signals for “bystander” genes (those not directly implicated in disease pathogenesis) may be easier to detect than those actually mediating the signal.

Finally, it is critical to emphasize that, even when co-localization has been demonstrated between a GWAS variant and a tissue-appropriate eQTL signal, this does constitute proof that the eGene concerned mediates disease predisposition. Causal relationships other than “variant to gene to disease” are possible, including the possibility the variant has horizontally pleiotropic effects on each^57^. Growing understanding of the extent of shared local regulatory activity and regulatory pleiotropy makes such an alternative explanation all the more credible^58^. It is best to regard genes highlighted by coincident GWAS and eQTL signals as “candidate” effector transcripts, and to proceed to experimental approaches that enable direct tests of causality. These may involve perturbing the gene across a range of disease-relevant cell-lines and animal models, and determining the impact on phenotypic readouts that represent reliable surrogates of disease pathophysiology.

## Methods

### Pancreatic Islet sample collection and processing

#### Geneva samples

Islet sample procurement, mRNA processing and sequencing procedure has been described in Nica et al.^17^. Briefly, Islets isolated from cadaveric pancreas were provided by the Cell Isolation and Transplant Center, Department of Surgery, Geneva University Hospital (Drs. T. Berney and D. Bosco) through the Juvenile Diabetes Research Foundation (JDRF) award 31-2008-416 (ECIT Islet for Basic Research Program). mRNA was extracted using RLT buffer (RNeasy, Qiagen) and total RNA was prepared according to the standard RNeasy protocol. The original RNA libraries were 49-bp paired-end sequenced however, in order to allow joint analysis with the other available datasets for this study, mRNA samples were re-processed using a 100-bp paired-end sequencing protocol. The library preparation and sequencing followed customary Illumina TruSeq protocols for next generation sequencing as described in the original publication^17^. All procedures followed ethical guidelines at the University Hospital in Geneva.

#### Lund Samples

Islet sample procurement, mRNA processing and sequencing procedure has been described in Fadista et al.^15^. Along with the 89 islet samples previously published^15^, we included 102 islet samples and processed these uniformly following the same protocol. These islet samples were obtained from 191 cadaver donors of European ancestry from the Nordic Islet Transplantation Programme (http://www.nordicislets.org). Purity of islets was assessed by dithizone staining, while measurement of DNA content and estimate of the contribution of exocrine and endocrine tissue were assessed as previously described^59^. Total RNA was isolated with the AllPrep DNA/RNA Mini Kit following the manufacturer’s instructions (Qiagen), sample preparation was performed using Illumina’s TruSeq RNA Sample Preparation Kit according to manufacturer’s recommendations. The target insert size of 300 bp was sequenced using a paired end 101 bp protocol on the HiSeq2000 platform (Illumina). Illumina Casava v.1.8.2 software was used for base calling. All procedures were approved by the ethics committee at Lund University.

#### Oxford samples

Samples collected in Oxford and Edmonton that were jointly sequenced in Oxford are included in this set of samples. Islet sample procurement, mRNA processing and sequencing procedure has been described in van de Bunt et al.^16^. To the 117 samples previously published (78 from Edmonton and 39 from Oxford), 57 samples were added and processed following similar protocols as before (27 from Edmonton and 30 from Oxford). Briefly, freshly isolated human islets were collected at the Oxford Centre for Islet Transplantation (OXCIT) in Oxford, or the Alberta Diabetes Institute IsletCore (www.isletcore.ca) in Edmonton, Canada. Additional islets were obtained from the Alberta Diabetes Institute IsletCore’s long-term cryopreserved biobank. Freshly isolated islets were processed for RNA and DNA extraction after 1–3 days in culture in CMRL media. Cryopreserved samples were thawed as described in Manning et al.^60^ and Lyon at al.^61^. RNA was extracted from human islets using Trizol (Ambion, UK or Sigma Aldrich, Canada). To clean remaining media from the islets, samples were washed three times with phosphate buffered saline (Sigma Aldrich, UK). After the final cleaning step 1 mL Trizol was added to the cells. The cells were lysed by pipetting immediately to ensure rapid inhibition of RNase activity and incubated at room temperature for ten minutes. Lysates were then transferred to clean 1.5 mL RNase-free centrifuge tubes (Applied Biosystems, UK). RNA quality (RIN score) was determined using an Agilent 2100 Bioanalyser (Agilent, UK), with a RIN score > 6 deemed acceptable for inclusion in the study. Samples were stored at −80°C prior to sequencing. Poly-A selected libraries were prepared from total RNA at the Oxford Genomics Centre using NEBNext ultra directional RNA library prep kit for Illumina with custom 8bp indexes^62^. Libraries were multiplexed (3 samples per lane), clustered using TruSeq PE Cluster Kit v3, and paired-end sequenced (100nt) using Illumina TruSeq v3 chemistry on the Illumina HiSeq2000 platform. All procedures were approved by the Human Research Ethics Board at the University of Alberta (Pro00013094), the University of Oxford’s Oxford Tropical Research Ethics Committee (OxTREC Reference: 2–15), or the Oxfordshire Regional Ethics Committee B (REC reference: 09/H0605/2). All organ donors provided informed consent for use of pancreatic tissue in research.

#### USA samples

Islet sample procurement, mRNA processing and sequencing has been described in Varshney et al.^14^. Briefly, 39 Islet samples from organ donors were received from the Integrated Islet Distribution Program, the National Disease Research Interchange (NDRI), and Prodo-Labs. Total RNA from 2000-3000 islet equivalents (IEQ) was extracted and purified using Trizol (Life Technologies). RNA quality was confirmed with Bioanalyzer 2100 (Agilent); samples with RNA integrity number (RIN) > 6.5 were prepared for mRNA sequencing. We added External RNA Control Consortium (ERCC) spike-in controls (Life Technologies) to one microgram of total RNA. PolyA+, stranded mRNA RNA-sequencing libraries were generated for each islet using the TruSeq stranded mRNA kit according to manufacturer’s protocol (Illumina). Each islet RNA-seq library was barcoded, pooled into 12-sample batches, and sequenced over multiple lanes of HiSeq 2000 to obtain an average depth of 100 million 2 × 101 bp sequences. All procedures followed ethical guidelines at the National Institutes of Health (NIH.)

### Beta-cell sample collection and processing

Sample collection, mRNA processing and sequencing procedure has been described in Nica et al.^17^. To the 11 FAC sorted beta-cells population samples previously published, we added 15 more samples that were processed following the same protocols. Briefly, islets were dispersed into single cells, stained with Newport Green, and sorted into ‘‘beta’’ and ‘‘non-beta’’ populations as described previously^63^. The proportion of beta (insulin), alpha (glucagon), and delta (somatostatin) cells in each population (as percentage of total cells) was determined by immunofluorescence. mRNA extractions as well as sequencing followed the same details described for islets samples processing for the Geneva samples.

### Read-mapping and exon quantification

The 100-bp sequenced paired-end reads were mapped to the GRCh37 reference genome^64^ with GEM ^65^. Exon quantifications were calculated for all elements annotated in GENCODE^66^ v19, removing genes with more than 20% zero read count. All overlapping exons of a gene were merged into meta-exons with identifier of type ENSG000001.1_exon.start.pos_exon.end.pos, as described in Lappalainen^20^. Read counts over these elements were calculated without using read pair information, except for excluding reads where the pairs mapped to two different genes. We counted a read in an exon if either its start or end coordinates overlapped an exon. For split reads, we counted the exon overlap of each split fragment, and added counts per read as 1/(number of overlapping exons per gene). Gene level quantifications used the sum of all reads mapped to exons from the gene. Genes with more than 20% zero read counts were removed.

### Genotype imputation

Genotypes for all islet samples, including 19 beta-cell samples, were available from omniexpress and omni2.5 genotype arrays. Quality of genotyping from the shared SNPs in both arrays was assessed before imputation separately by removing SNPs as follows: 1) SNPs with minor allele frequency (MAF) < 5%; 2) SNP genotype success rate <95%; 3) Palindromic SNPs with MAF > 40%; 4) HWE < 1e-6; 5) Absence from 1000G reference panel; 6) Allele inconsistencies with 1000G reference panel; 7) Probes for same rsID mapping to multiple genomic locations (1000G reference-consistent probe kept). Finally, samples were excluded if they had an overlap genotype success rate lower than 90%; and MAF differences larger than 20% compared to the 1000G reported european MAF.

The two panels were separately pre-phased with SHAPEIT^67^ v2 using the IMPUTE2-supplied genetic maps. After pre-phasing the panels were imputed with IMPUTE2^68^ v2.3.1 using the 1000 Genomes Phase I integrated variant set (March 2012) as the reference panel^69^. SNPs with INFO score > 0.4 and HWE p > 1e-6 (for chrX this was calculated from female individuals only) from each panel were kept. A combined vcf for each chromosome was generated from the intersection of the checked variants in each panel. Directly genotyped SNPs with a MAF < 1% (including the exome-components of the chips not shared between all centres) were merged into the combined vcfs: i) If SNPs were not imputed they were added and ii) If SNPs had been imputed, the imputed calls for the individual were replaced by the typed genotype. Dosages were calculated from the imputation probabilities (genotyped samples) or genotype calls (WGS samples). For the 22 autosomes the dosage calculation was: 2× ((0.5*heterozygous call) + homozygous alt call). For chromosome X (where every individual should be functionally hemizygous), the calculation was: (0.5*heterozygous call) + homozygous alt call). Genotype calls for males can only be ‘0/0’ and ‘1/1’. The total number of variants available for analysis after quality assessment was 8,056,952.

For the 26 beta cell samples, 19 had genotypes available from omniexpress arrays, whereas 7 had the DNA sequence available. Variant calling from DNA sequence has been previously described in Nica et al.^17^. Briefly, the Genome Analysis Toolkit (GATK)^70^ v1.5.31 was used following the Best Practice Variant Detection v3 to call variants. Reads were aligned to the hg19 reference genome with BWA^71^. We used a confidence score threshold of 30 for variant detection and a minimum base quality of 17 for base calling. Good confidence (1% FDR) SNP calls were then imputed on the 1000 Genomes reference panel and phased with BEAGLE^72^ v3.3.2. Imputation of variants from samples with arrays genotyping were imputed together with genotypes from individuals with islets samples as described before and then merged with genotypes from DNA sequences. SNPs with INFO score > 0.4, HWE p > 1e-6 and MAF > 5%, were kept for further analysis. The total number of variants available for analysis after quality assessment was 6,847,993.

### RNAseq quality assessment and data normalization

Heterozygous sites per sample were matched with genotype information to confirm the ID of the samples^73^. 11 samples did not match with their genotypes, 6 of which would be corrected by identifying a good match. For the remaining samples, no matches were found on the genotypes and they were removed from the dataset, giving a total of 420 samples with genotypes.

Raw read counts from exons and genes were scaled to 10 million to allow comparison between samples with different libraries. Scaled raw counts were then quantile normalized. We used principal component analysis (PCA) to evaluate the effects of unwanted technical variation and the expected batch effects due to fact that the islet sample processing mRNA sequencing was performed across four labs. We evaluated a) the optimal number of principal components (PCs) for the discovery of eQTLs and b) the minimum number of PCs necessary to control for laboratories of origin batch effects (Supp. Figure SF1). We performed eQTL discovery controlling for 1,5,10, 20 30 40 and 50 PCs for expression, as well as gender, 4 PCs derived from genotype data, and a variable defining the laboratory of origin (coded as: OXF, LUND, GEN and USA). After evaluation of the results, we conclude that controlling for 20 PCs was optimal. To ensure that we controlled for batch effects with these variables, we used a permutation scheme as follows: expression sample labels and expression covariates were permuted within each of the 4 laboratories before performing a standard eQTL analysis against non-permuted genotypes (and matched PCs for genotypes) using different numbers of PCs for expression. Significant eQTLs in any of these analyses are considered a false positive due to technical differences across laboratories of origin of the samples. Our results indicate that 10PCs were sufficient to minimize the number of false positives due to batch effects originating from differences in processing of the islet samples (Supp. Figure SF1).

### eQTL analysis

eQTL analysis for islets and beta-cells were performed using fastQTL^18^ on 420 islets samples and 26 beta-cells samples with available genotypes. Cis-eQTL analysis was restricted to SNPs in a 1MB window upstream and downstream the transcription start site (TSS) for each gene and SNPs with MAF>1%. For the analysis of beta-cell samples, we used a filter of MAF>5%. Exon-level eQTLs identified best exons-SNP association per gene (using the –group flag), while gene level eQTLs used gene quantifications and identified the best gene-SNP association. Variables included in the linear models were the first 4 PCs for genotypes, the first 25 PCs for expression, gender and a variable identifying the laboratory of origin of the samples. Significance for the SNP-gene association was assessed using 1000 permutations per gene, correcting P values with a beta approximation distribution^18^. Genome-wide multiple testing correction was performed using the q-value correction^21^ implemented in largeQvalue^74^.

Results of this joint analysis were highly-correlated with those obtained from a fixed-effects meta-analysis of the four component studies, indicating appropriate control for the technical differences between the studies (Supp Figure SF14).

To discover multiple independent eQTLs, we applied a stepwise regression procedure as described in Brown et al.^75^. Briefly, we started from the set of eGenes discovered in the first pass of association analysis (FDR < 1%). Then, the maximum beta-adjusted P value (correcting for multiple testing across the SNPs and exons) over these genes was taken as the gene-level threshold. The next stage proceeded iteratively for each gene and threshold. A cis-scan of the window was performed in each iteration, using 1,000 permutations and correcting for all previously discovered SNPs. If the beta adjusted P value for the most significant exon-SNP or gene-SNP (best association) was not significant at the gene-level threshold, the forward stage was complete and the procedure moved on to the backward step. If this P value was significant, the best association was added to the list of discovered eQTLs as an independent signal and the forward step proceeded to the next iteration. The exon level cis-eQTL scan identified eQTLs from different exons, but reported only the best exon-SNP in each iteration. Once the forward stage was complete for a given gene, a list of associated SNPs was produced which we refer to as forward signals. The backward stage consisted of testing each forward signal separately, controlling for all other discovered signals. To do this, for each forward signal we ran a cis scan over all variants in the window using fastQTL, fitting all other discovered signals as covariates. If no SNP was significant at the gene-level threshold the signal being tested was dropped, otherwise the best association from the scan was chosen as the variant that represented the signal best in the full model.

The principal component adjustments used to control for unwanted technical variation during eQTL analysis were designed to account for differences in sample processing across laboratories and for some of the impact of variation in purity between samples. However, by correlating the data-generated PCs with cell proportion estimates, we observed that, even when adjusting using 25 PCs, only ∼30% of the variance attributable to variation in exocrine or beta-cell composition was regressed out, and that more than 100 PCs were required to remove at least 50% of the variance. This indicates that some of the eQTLs here attributed to pancreatic islets may, in fact, reflect exocrine pancreatic contamination. To evaluate this further, we compared the sets of eQTLs identified in the InsPIRE islet samples with the highest and lowest proportions of exocrine contamination (n=100 for each) and 100 randomly-selected GTEx whole pancreas samples. Overlap between whole pancreas and islet eQTLs was greater in islet samples with the highest exocrine contamination (π_1_ 75% vs 64%), indicating that cell-specific effects were preserved even controlling for 25 PCs for the eQTL analysis (Supp. Figure SF15).

### GTEx eQTLs

We identified exon level eQTLs for 44 GTEx tissues using fastQTL^18^ following the same procedure as for the islet eQTLs. Covariates included followed the previously published number of PCs for expression^7^ and included 15 PCs for expression for tissues with less than 154 samples; 30 PCs for samples between 155 and 254 samples; and 35 PCs for samples with more than 254 samples. Independent eQTLs from exons were identified as described for islets eQTLs. The proportion of shared eQTLs between islet and beta-cell eQTLs and the eQTLs from GTEx tissues were identified using Π1^21^.

### Tissue de-convolution

To identify the contribution of the beta-cells, non-beta cells and exocrine components (non-islets cell) expression to the total gene expression measure in islets we performed an expression deconvolution analysis. Expression profiles from GTEx whole pancreas was used as a model for the exocrine component of expression^7^, while FAC-sorted expression profiles from beta-cell and non-beta-cells from Nica et al.^17^ were used to identify the fraction of expression derived from islets cells. First, we performed differential expression analysis of a) exocrine versus whole islet samples; b) beta-cell versus whole islet samples; c) non-beta-cell versus whole islet samples. The top 500 genes from each analysis were combined, and a deconvolution matrix of log_2_-transformed median expression values was prepared for each cell type. Next, deconvolution was performed using the Bioconductor package DeconRNASeq^76^. Deconvolution values per sample are included in the covariates file, together with the expression values in the EGA submission.

### Enrichment of eQTLs in T2D and glycemic GWAS

To perform an enrichment analysis of T2D and glycemic traits GWAS associations among eQTLs across tissues, we examined 78 T2D associated signals^3^, and 44 variants from associations with continuous glycemic traits relevant to T2D predisposition (including fasting glucose, and beta-cell function (HOMA-B) in non-diabetic individuals) (Supp. Table ST13)^34,36,60^. For each GWAS lead variant, we extracted the eQTL with the greatest absolute effect size estimate from the results for all GTEx tissues and the InsPIRE pancreatic islets. We then compared their observed effect size estimates to those derived from a null distribution of 15,000 random variants matched in terms of the number of SNPs in LD, distance to TSS, number of nearby genes and minor allele frequency. For comparison with results observed for T2D loci, we also included the set of 50 lead variants implicated by GWAS in predisposition to T1D^37^.

### Co-localization of islet eQTL with T2D GWAS

Co-localization of GWAS variants and eQTLs were performed using both COLOC^40^ and RTC^41^. For the analysis using COLOC, all variants within 250 kilobase flanking regions around the index variants were tested for co-localization using default parameters from the software were used on summary statistics from T2D GWAS^5^ and fasting glucose^35^. GWAS variants and eSNPs pairs were considered to co-localize if the COLOC score for shared signal was larger than 0.9. RTC analysis was also performed using defaults parameters from the software with a list of 86 lead GWAS variants for T2D and fasting glucose (Supp. Table ST13). Associations between GWAS and gene expression were considered as co-localizing if RTC score was larger than 0.9 (Supp. Table ST15).

### Chromatin states, Islet ATAC-seq and Transcription factor (TF) footprints

We used a previously published 13 chromatin state model that included Pancreatic Islets along with 30 other diverse tissues^14^. Briefly, these chromatin states were generated from cell/tissue ChIP-seq data for H3K27ac, H3K27me3, H3K36me3, H3K4me1, and H3K4me3, and input control from a diverse set of publically available data^29,77-79^ using the ChromHMM program^80^. Chromatin states were learned jointly from 33 cell/tissues that passed QC by applying the ChromHMM (version 1.10) hidden Markov model algorithm at 200-bp resolution to five chromatin marks and input^14^. We ran ChromHMM with a range of possible states and selected a 13-state model, because it most accurately captured information from higher-state models and provided sufficient resolution to identify biologically meaningful patterns in a reproducible way. As reported previously^14^, *Stretch Enhancers* were defined as contiguous enhancer chromatin state (Active Enhancer 1 and 2, Genic Enhancer and Weak Enhancer) segments longer than 3kb, whereas *Typical Enhancers* were enhancer state segments smaller than the median length of 800bp^29^.

We used the union of ATAC-seq peaks previously identified from two human islet samples called using MACS2 v2.1.0 (https://github.com/taoliu/MACS). We also used previously published TF footprints that were generated in a haplotype-aware manner using ATAC-seq and genotyping data from the phased, imputed genotypes for each of two islet samples using vcf2diploid^81^ v0.2.6a.

### Filtering eQTL SNPs for epigenomic analyses

Since low MAF SNPs, due to low power, can only be identified as significant eQTL SNP (eSNPs) with high eQTL effect sizes (slope or the beta from the linear regression), we observed that absolute effect size varies inversely with MAF (Supp. Figure SF13). To conduct eQTL effect size based analyses in an unbiased manner, we selected significant (FDR 1%) eSNPs with MAF>=0.2. We then pruned this list to retain the most significant SNPs with pairwise LD(r2)<0.8 for the EUR population using PLINK^82^ and 1000 genomes variant call format (vcf) files (downloaded from ftp://ftp.1000genomes.ebi.ac.uk/vol1/ftp/release/20130502/) for reference (European population). This filtering process resulted in n=3832 islet eSNPs.

### Enrichment of genetic variants in genomic features

To calculate the enrichment of islet eSNPs to overlap with genomic features such as chromatin states and transcription factor (TF) footprint motifs, we used the GREGOR tool^83^. For each input SNP, GREGOR selects ∼500 control SNPs matched for MAF, distance to the gene, and number of SNPs in LD(r2)≥0.99. A unique overlap is reported if the feature overlaps any input lead SNP or its LD(r2)>0.99 LD SNPs. Fold enrichment is calculated as the number unique overlaps over the mean number of loci at which the matched control SNPs (or their LD(r2>)0.99 SNPs) overlap the same feature. This process accounts for the length of the features, as longer features will have more overlap by chance with control SNP sets. We used the following parameters in GREGOR for eQTL enrichment: r2 threshold (for inclusion of SNPs in linkage disequilibrium (LD) with the lead eSNP)=0.99, LD window size=1Mb, and minimum neighbor number=500.

For enrichment of T2D GWAS SNPs in islet chromatin states, we downloaded the list of T2D GWAS SNPs from Mahajan, et al.^33^. We pruned this list to retain the most significant SNPs with pairwise LD(r2)<0.2 for the EUR population using PLINK^82^ and 1000 genomes variant call format (vcf) files (downloaded from ftp://ftp.1000genomes.ebi.ac.uk/vol1/ftp/release/20130502/) for reference (European population). This filtering process resulted in N=378 T2D GWAS SNPs. We used GREGOR to calculate enrichment using the following specific parameters: r2 threshold (for inclusion of SNPs in linkage disequilibrium (LD) with the lead eSNP)=0.8, LD window size=1Mb, and minimum neighbor number=500.

We investigated if footprint motifs were more enriched to overlap eQTL of high vs low effect sizes. We sorted the filtered (as described above) eQTL list by absolute effect size values and partitioned into two equally sized bins (N eSNPs=1,916). Since TF footprints were available for a large number of motifs (N motifs=1,995), the enrichment analysis had a large multiple testing burden and limited power with 1,916 eSNPs in each bin. Therefore, we only considered footprint motifs that were significantly enriched (FDR<1%, Benjamini & Yekutieli method from R p.adjust function, N motifs=283) to overlap the bulk set of eSNPs (LD r2<0.8 pruned but not MAF filtered, N eSNPs=6,468, Supp. Table ST10) for enrichment to overlap the binned set of eSNPs. This helped reduce the multiple testing burden. We then calculated enrichment for the selected footprints to overlap SNPs in each bin using GREGOR with same parameters as described above (Supp. Table ST11).

### eSNP effect size distribution in chromatin states and ATAC-seq peaks within chromatin states

We identified the islet eQTL eSNPs (after LD pruning and MAF filtering as described above) occurring in chromatin states or ATAC-seq peaks within chromatin states using BEDtools intersect^84^. Similar to the enrichment calculation procedure, we considered a unique eQTL overlap if the lead eSNP or a proxy SNP with LD(r^2^)>0.99 occurred in these regions. We considered the effect size as the slope or the beta from the linear regression for the eQTL overlapping each region. *P* values were calculated using the Wilcoxon Rank Sum Test in R^85^.

### TF motif directionality analysis

For TF footprint motifs that were significantly enriched to overlap the full set of islet eQTLs (after LD pruning to r2<0.8) with (FDR 1%, Benjamini & Yekutieli method from R p.adjust function, N motifs= 283), we determined the overlap position of the eSNP (pruned LD r2<0.8 lead eSNPs and their LD r2>0.99 proxy SNPs) with each TF footprint motif. We considered instances where the eSNP overlapped the TF footprint motif at a position with information content >=0.7 and either the eSNP effect or the non-effect allele was the most preferred base in the motif. We selected TF footprint motifs that had 10 or more such eSNP overlap instances (N=278). For each TF footprint motif and eSNP overlap, we re-keyed the direction of effect on the target gene being positive or negative with respect to the most preferred base in the motif. For each TF motif, we compiled the fraction of instances where the SNP allele that was most preferred in the TF footprint motif (i.e. base with highest probability in the motif) associated with increased expression of the associated gene. We refer to this metric as the motif directionality fraction where fraction near 1 suggests activating and fraction near 0 suggests repressive preferences towards the target gene expression. Motif directionality fraction near 0.5 suggests no activity preference or context dependence.

We compared our results to a previously published study that quantified transcription activating or repressive activities based on massively parallel reported assays in HepG2 and K562 cells^32^ (Supp. Figure SF8). We found that the motif directionality measures metric were largely concordant (Spearman’s r=0.64, *P*=8.1×10^-13^) with orthogonal motif activity measures derived from massively parallel reporter assays (MPRAs) performed in HepG2 and K562 cell lines^32^ (Supp. Figure SF8). We then considered 99 motifs from our analyses that were reported to have significant (P<0.01) activating or repressive scores from MPRAs in both HepG2 and K562. With the null expectation of the motif directionality fraction being equal to 0.5, i.e. TF binding equally likely to increase or decrease target gene expression, we used a binomial test to calculate TF that show significant deviation from the null (N=8 at FDR < 10%, Supp. Table ST11).

### Cell culture

MIN6 mouse insulinoma beta cells^50^ were grown in Dulbecco’s modified Eagle’s Medium (Sigma-Aldrich, St. Louis, Missouri/USA) with 10% fetal bovine serum, 1 mM sodium pyruvate, and 0.1 mM beta-mercaptoethanol. INS-1-derived 832/13 rat insulinoma beta cells (a gift from C. Newgard, Duke University, Durham, North Carolina/USA) were grown in RPMI-1640 medium (Corning, New York/USA) supplemented with 10% fetal bovine serum, 10 mM HEPES, 2 mM L-glutamine, 1 mM sodium pyruvate, and 0.05 mM beta-mercaptoethanol. EndoC-βH1 cells (Endocell) were grown according to Ravassard et al.^49^ in Dulbecco’s modified Eagle’s medium (DMEM; Sigma-Aldrich), 5.6mmol/L glucose with 2% BSA fraction V fatty acid free (Roche Diagnostics), 50μmol/L 2-mercaptoethanol, 10mmol/L nicotinamide (Calbiochem), 5.5μg/ml transferrin (Sigma-Aldrich), 6.7ng/ml selenite (Sigma-Aldrich), 100U/ml penicillin, and 100μg/ml streptomycin. Cells were grown on coating consisting of 1% matrigel and 2µg/mL fibronectin (Sigma). We maintained cell lines at 37° C and 5% CO2.

### Transcriptional reporter assays

To test haplotypes for allele-specific effects on transcriptional activity, we PCR-amplified a 765-bp genomic region (haplotype A) containing variants: rs7798124, rs7798360, and rs7781710, and a second 592-bp genomic region (haplotype B) containing variants: rs10228796, rs10258074, rs2191348, and rs2191349 from DNA of individuals homozygous for each haplotype. The oligonucleotide primer sequences are listed in Supp. Table ST17. We cloned the PCR amplicons into the multiple cloning site of the Firefly luciferase reporter vector pGL4.23 (Promega, Fitchburg, Wisconsin/USA) in both orientations, as described previously^86^. Vectors are designated as ‘forward’ or ‘reverse’ based on the PCR-amplicon orientation with respect to DGKB gene. We isolated and verified the sequence of five independent clones for each haplotype in each orientation. For the 5’ eQTL a 250 bp construct containing the rs17168486 SNP (Origene) was subcloned into the Firefly luciferase reporter vector pGL4.23 (Promega) in both orientations.

We plated the MIN6 (200,000 cells) or 832/13 (300,000 cells) in 24-well plates 24 hrs before transfections and the EndoC-βH1 cells (140.000 cells) plated 48H prior to transfection. We co-transfected the pGL4.23 constructs with phRL-TK Renilla luciferase reporter vector (Promega) in duplicate into MIN6 or 832/13 cells and in triplicate for EndoC-βH1 cells. For the transfections we used Lipofectamine LTX (ThermoFisher Scientific, Waltham, Massachusetts/USA) with 250 ng of plasmid DNA and 80 ng Renilla for MIN6 cells, Fugene6 (Promega) with 720 ng of plasmid and 80 ng Renilla for 832/13 cells per each welll and Fugene6 with 700 ng plasmid and 10 ng renilla for EndoC-βH1 cells. We incubated the transfected cells at 37° C with 5% CO2 for 48 hours. We measured the luciferase activity with cell lysates using the Dual-Luciferase® Reporter Assay System (Promega). We normalized Firefly luciferase activity to the Renilla luciferase activity. We compared differences between the haplotypes using unpaired two-sided t-tests. All experiments were independently repeated on a second day and yielded comparable results.

### Electrophoretic Mobility Shift Assays

Electrophoretic mobility shift assays were performed as previously described. We annealed 17-nucleotide biotinylated complementary oligonucleotides (Integrated DNA Technologies) centered on variants: rs10228796, rs10258074, rs2191348, and rs2191349 (Supp. Table ST18). MIN6 nuclear protein extract was prepared using the NE-PER kit (Thermo Scientific). To conduct the EMSA binding reactions, we used the LightShift Chemiluminescent EMSA kit (Thermo Scientific) following the manufacturer’s protocol. Each reaction consisted of 1 μg poly(dI-dC), 1× binding buffer, 10 μg MIN6 nuclear extract, 400 fmol biotinylated oligonucleotide. We resolved DNA-protein complexes on nondenaturing DNA retardation gels (Invitrogen) in 0.5×TBE. We transferred the complexes to Biodyne B Nylon membranes (Pall Corporation), and UV cross-linked (Stratagene) to the membrane. We used chemiluminescence to detect the DNA-protein complexes. EMSAs were repeated on a second day with comparable results.

## Acknowledgments/Funding

This work has been supported with grants awarded or supporting the following individuals: A. Viñuela and E.T. Dermitzakis were supported by EU IMI program (UE7-DIRECT-115317-1), NIH (NIH-R01MH101814) and FNS funded project RNA1 (31003A_149984). A. Varshney was supported by the American Association for University Women International Doctoral Fellowship, Barbour Doctoral Scholarship, and the University of Michigan Rackham Predoctoral Fellowship. MvdB was supported by a Novo Nordisk postdoctoral fellowship run in partnership with the University of Oxford. R. B. Prasad was supported by the EFSD/Novo Nordisk Programme for Diabetes Research in Europe, Diabetes Wellness (720-858-16 JDWG), Åke Wiberg Foundation (M18-0216). L. Groop was supported by the Swedish Research Council project grant (2015-2558) Swedish Research Council, AstraZeneca (10033731), Strategic Research Area Exodiab, Dnr 2009-1039, Swedish Foundation for Strategic Research Dnr IRC15-0067, and the Swedish Research Council, Linnaeus grant, Dnr 349-2006-237. F. S. Collins, M. Erdos, and N. Narisu were supported by NHGRI - ZIA HG000024. A. Iyengar, S. Vadlamudi and K. Mohlke were supported by NIH R01 DK072193, NIH U01 DK105561. S. C. J. Parker, L. J. Scott, and M. Boehnke were supported by U01DK062370. M. Stitzel was supported by K99/R00DK092251. P. Orchard was supported by grant T32 HG00040 from the National Human Genome Research Institute of the National Institutes of Health. P. MacDonald was supported by a Foundation grant from the Canadian Institutes of Health Research (CIHR: 148451). S. C. J. Parker was supported by National Institute of Diabetes and Digestive and Kidney Diseases grants R00 DK-099240 and R01 DK-117960, American Diabetes Association Pathway to Stop Diabetes grant 1–14-INI-7. The Alberta Diabetes Institute IsletCore was supported by the Alberta Diabetes Foundation. We thank the Human Organ Procurement and Exchange (HOPE) program and the Trillium Gift of Life Network (TGLN) for their efforts in obtaining human organs for research. A.L. Gloyn is a Wellcome Senior Fellow in Basic Biomedical Science. This work was funded in Oxford by the Wellcome Trust (095101, 200837, 106130, 203141, Medical Research Council (MR/L020149/1), European Union Horizon 2020 Programme (T2D Systems), and NIH (U01-DK105535; U01-DK085545). MMcC is a Wellcome Senior Investigator and an NIHR Senior Investigator. He was supported by the Wellcome Trust (Grants no. 090532, 106130, 098381, 203141, 212259); Medical Research Council grant no. MR/L020149/1; NIDDK (U01-DK105535, R01-MH101814, R01-MH090941); NIHR (NF-SI-0617-10090). This work was also supported by the Oxford NIHR Biomedical Research Centre. The views expressed in this article are those of the author(s) and not necessarily those of the NHS, the NIHR, or the Department of Health.

## Author Contributions

AVi, and CH performed the re-mapping, quantification and quality checks for the joint RNAseq dataset with assistance from MvdB, JF and NO. MvdB performed genotype quality evaluation and imputation of the joint genotypes data with assistance of AM. Geneva samples (GEN) were collected, processed and evaluated by AVi, CH, NIP, AAB and ETD. Lund samples (LUN) collected processed and evaluated by RBP, OA, JF, OH, GH, UK, NO, LG. Oxford and Edmonton samples (OXF) were collected processed and evaluated by MvdB, AB, PJ, PEM, AM, JEMF, VN, AP, ALG, MIM. USA samples as well as ATAC-seq were collected processed and evaluated by AVa, MB, MRE, NN, PO, MLS, RW, FSC, LJS, SCJP. Data analyses were performed by AVi, AVa, MvdB, RBP, AAB, JF, NO, AP, LJ. Experiments associated to the validation of DGKB eQTLs were performed by VN, SV, AKL, KLM, ALG, AVa and SCJP. The manuscript was drafted by AVi, AVa, MvdB, LJS, SCJP and MIM, then revised and approved by all authors.

## Supplemental Figures

**Supplemental Figure SF1.**
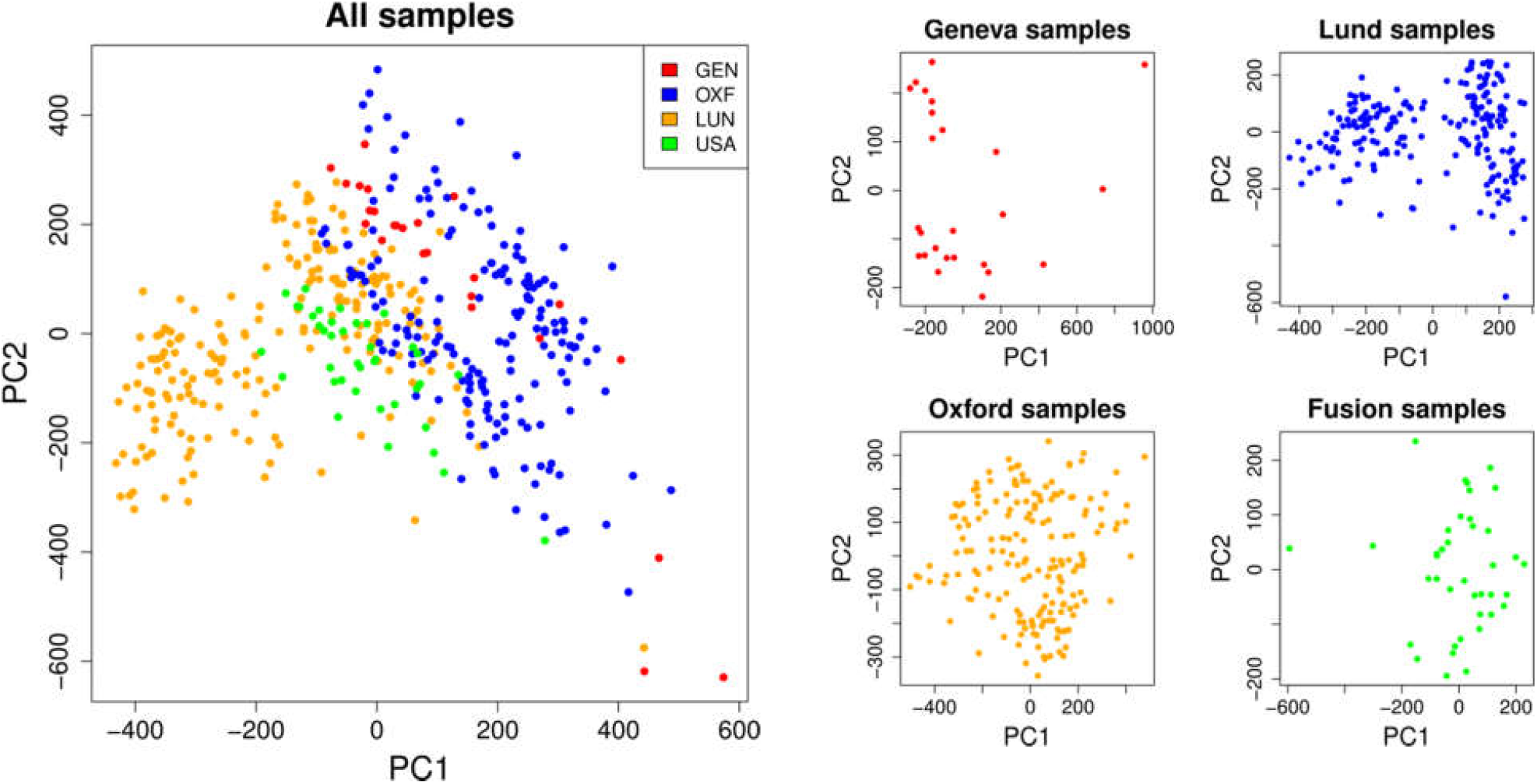
Principal component analysis (PCA) of the exon expression profiles per sample included in the InsPIRE project. Samples were re-quantified and normalized together to account for differences in the data production. The samples showed in the PCA analysis the differences due to experimental processing differences, with internal batch effects.

**Supplementary Figure SF2.**
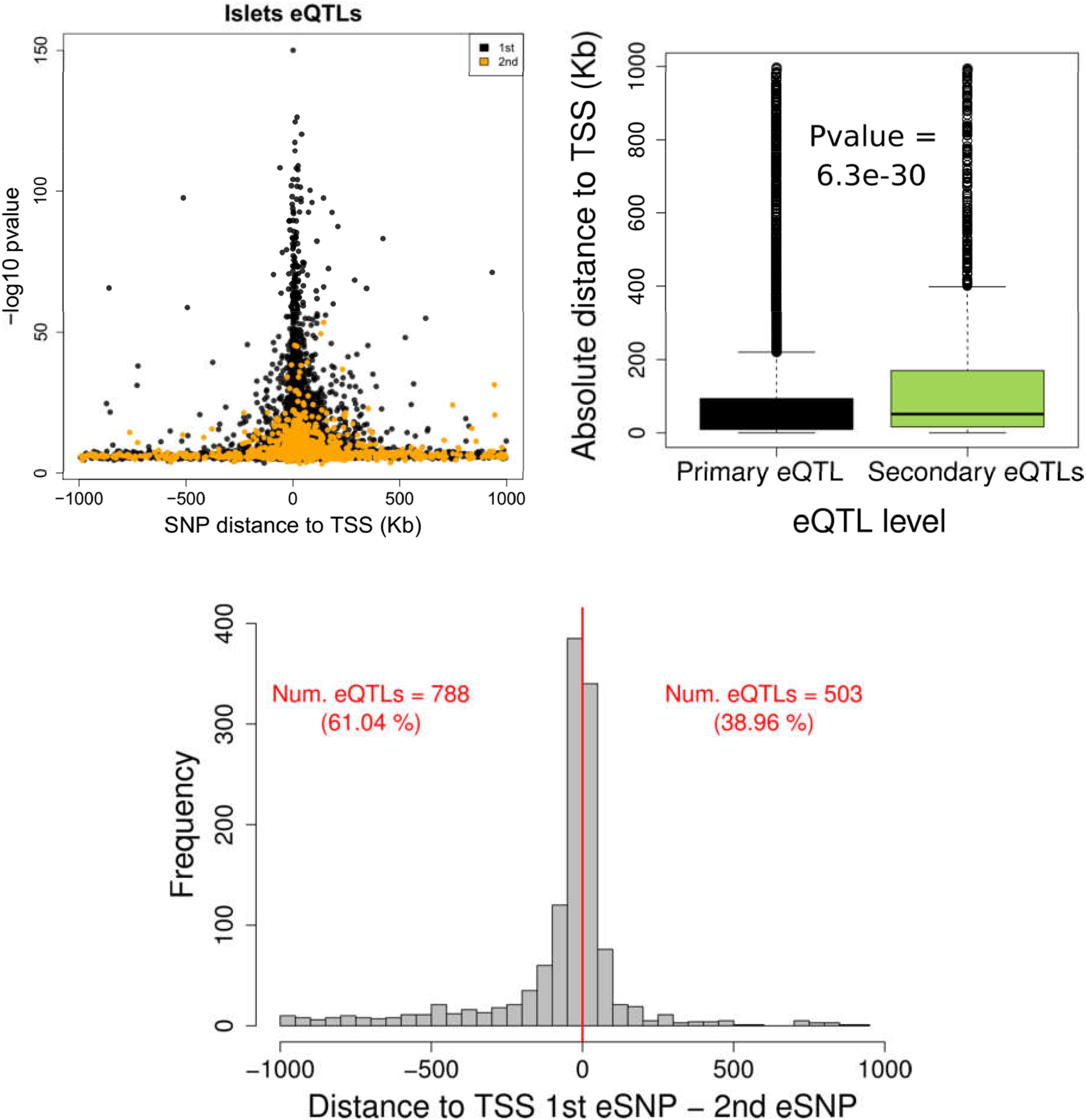
eQTL analysis. Top left figure shows the –log10 pvalue distribution of the lead eSNP per gene around the transcription start site (TSS) of the genes in black. Yellow values show the secondary signals discovered after conditional analysis. Both the primary and secondary sSNPs show smaller pvalues around the TSS, however, the secondary signals are significantly futher away from the TSS (top right plot). The bottom plot shows the distance of the eSNPs around the TSS for those genes with 2 indepdnetn eQTLs (n = 1,290). The difference in the Kb distance between primary SNP (1^st^) and secondary SNP (2nd SNP as the highest variance explained in expression) independent eSNPs significantly affecting the expression of the same gene is expressed in negative values (left) if the primary eSNPs is closer to the TSS than the secondary eSNPs (N = 788). Positive values identify those eGenes in which the secondary eSNP is closer to the TSS than the primary (N= 503).

**Supplementary Figure SF3.**
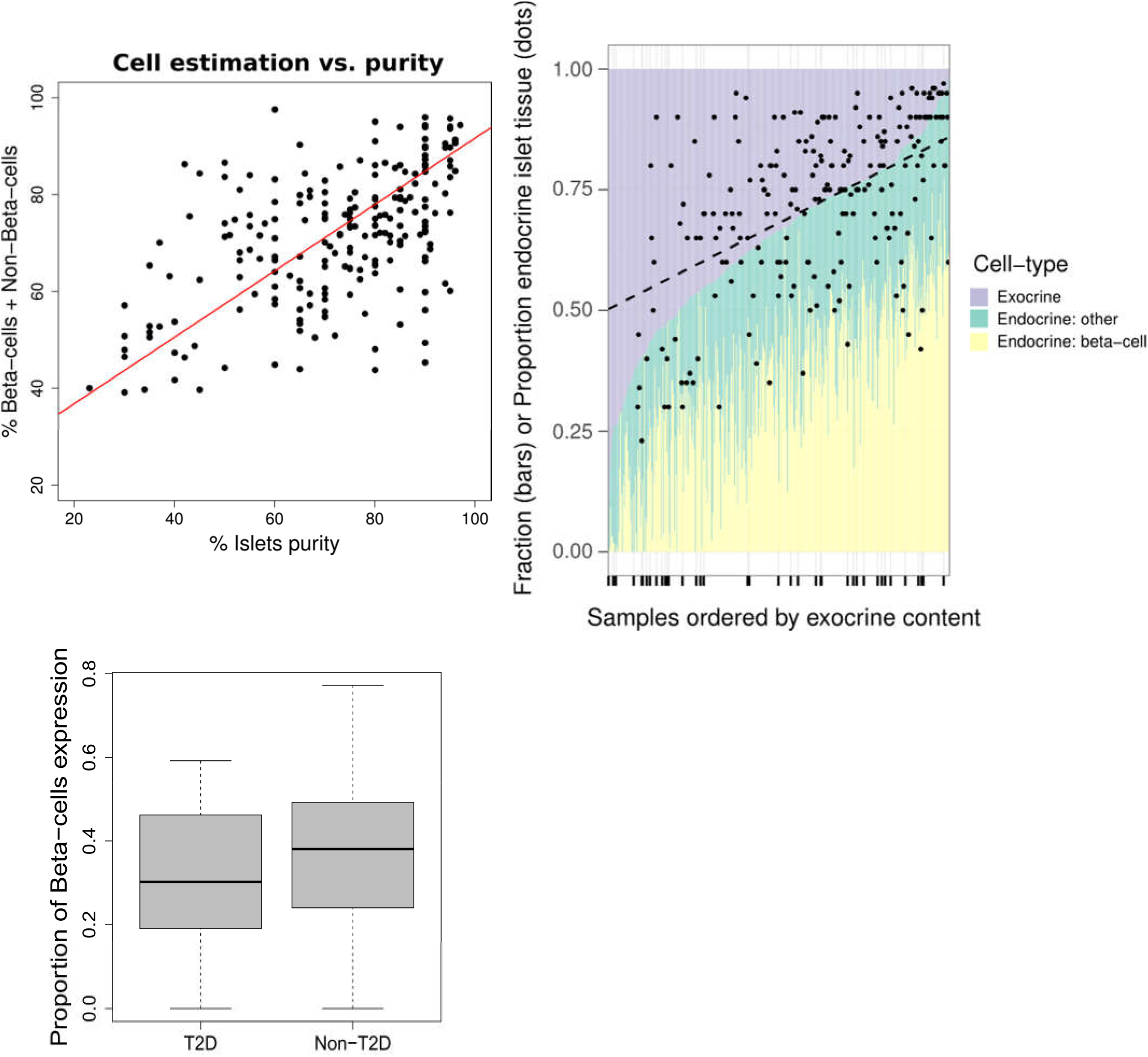
Cell deconvolution analysis. Top right plot shows the estimates of the different types of cell considered in the 420 islets samples processed. The beta-cells proportion composition form per sample corresponded to a median os 58.8%, and 41.2% for non-beta-cell fractions. Top left plot shows the percentage of purity for islets as measured in dithizone staining of the 232 samples compare to the estimated proportion of (beta-cells + other non-exocrine cell)/ total cell content in islets. The correlation between measured values of purity was ρ=-0.5 (P=2.8×10^-15^). Bottom plot shows the Percentage of Beta-cells expression detected in islets samples from individuals identified as diabetics (T2D), compare to non-T2D individuals.

**Supplementary Figure SF4.**
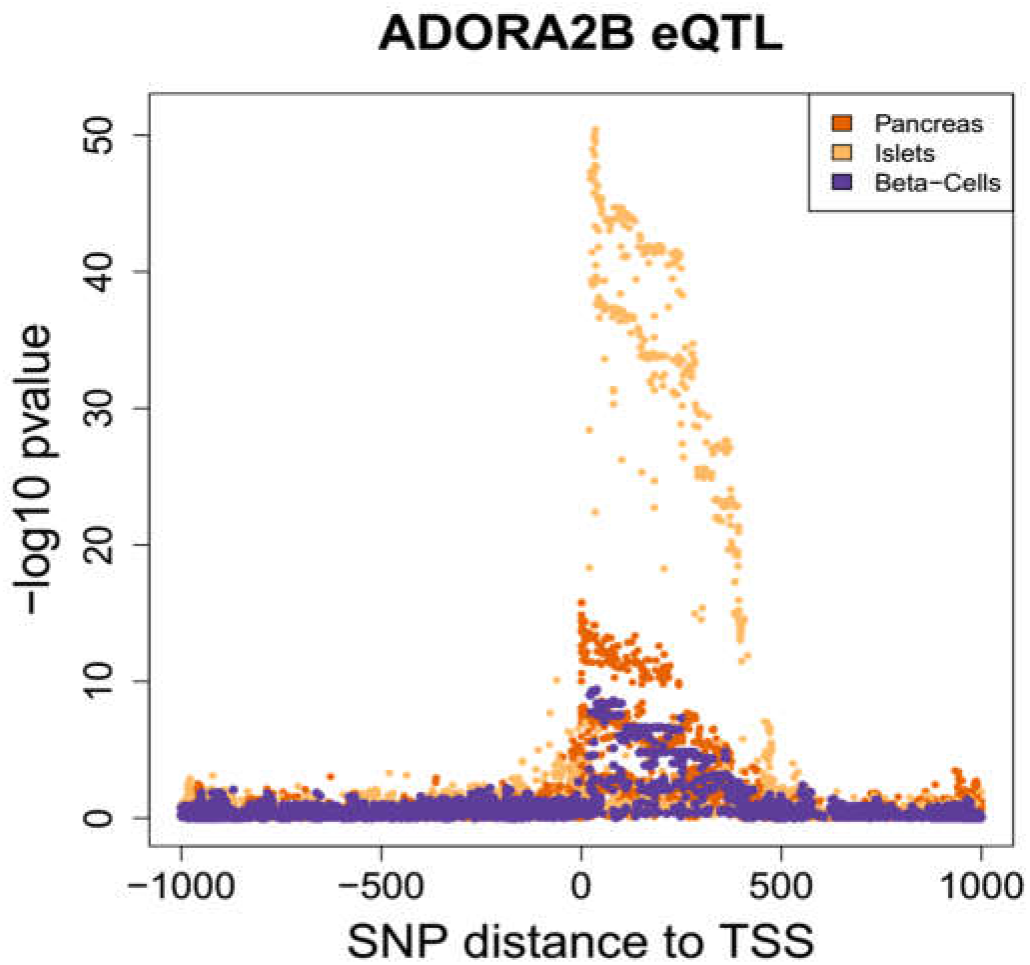
eQTL for ADORA2B in islets, beta-cells and pancreas samples. Each dot represent a SNPs in the cis window of ADORA2B and their distance in kb to the TSS. The y-axis shows the −log10 of the P value for the association between a given SNP and the expression of the same exon in ADORA2B (exon ID =ENSG00000170425.3_15877177_15879060). For all tissues, at least one SNP was significant after multiple testing (FDR = 5%).

**Supplemental Figure SF5.**
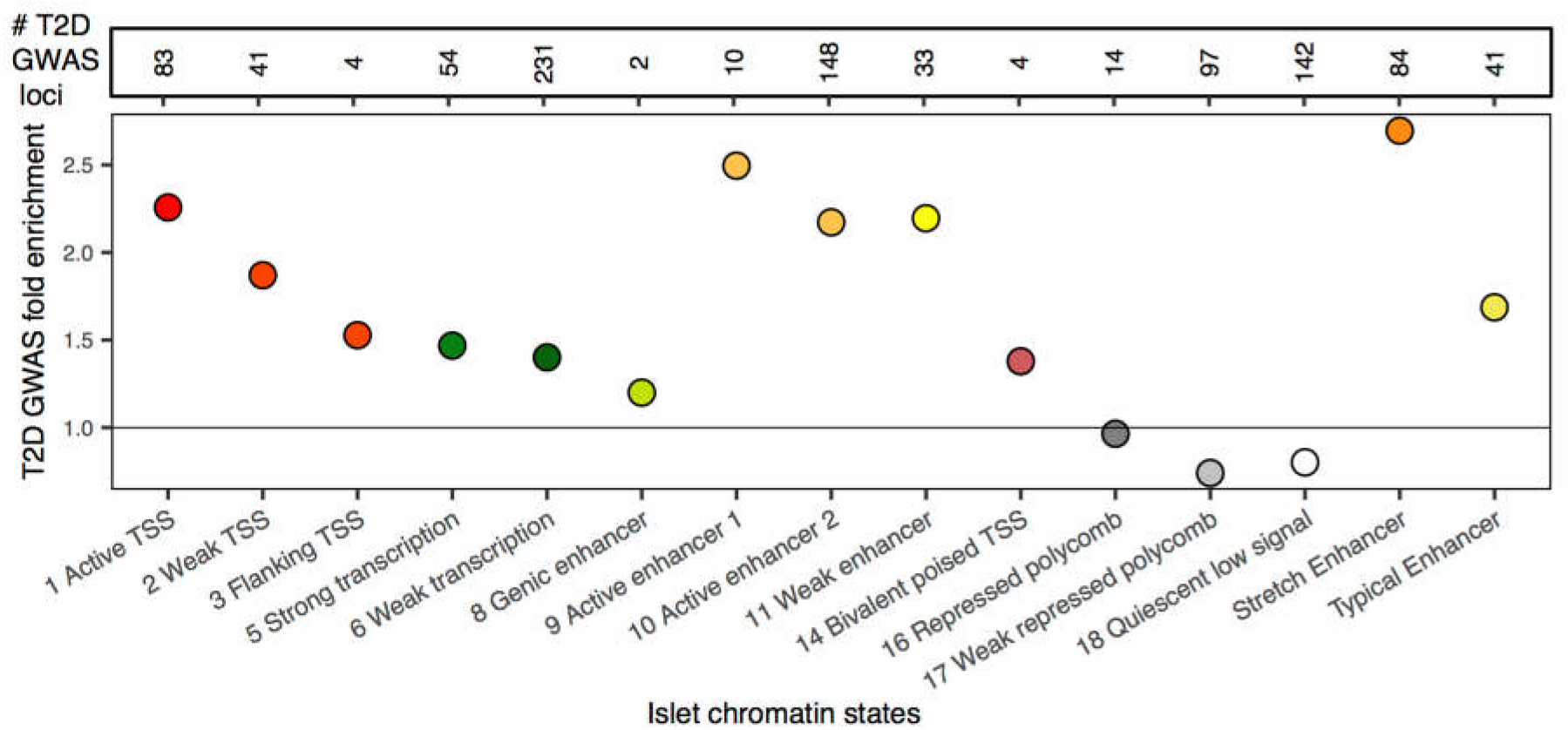
Islet eQTL overlap and enrichment in islet chromatin states. Top: Number of islet eQTL in 13 islet chromatin states and stretch and typical enhancers. Bottom: Fold enrichment of islet eQTL in chromatin states calculated using GREGOR (CITE: Schmidt 2015 Bioinformatics).

**Supplemental Figure SF6.**
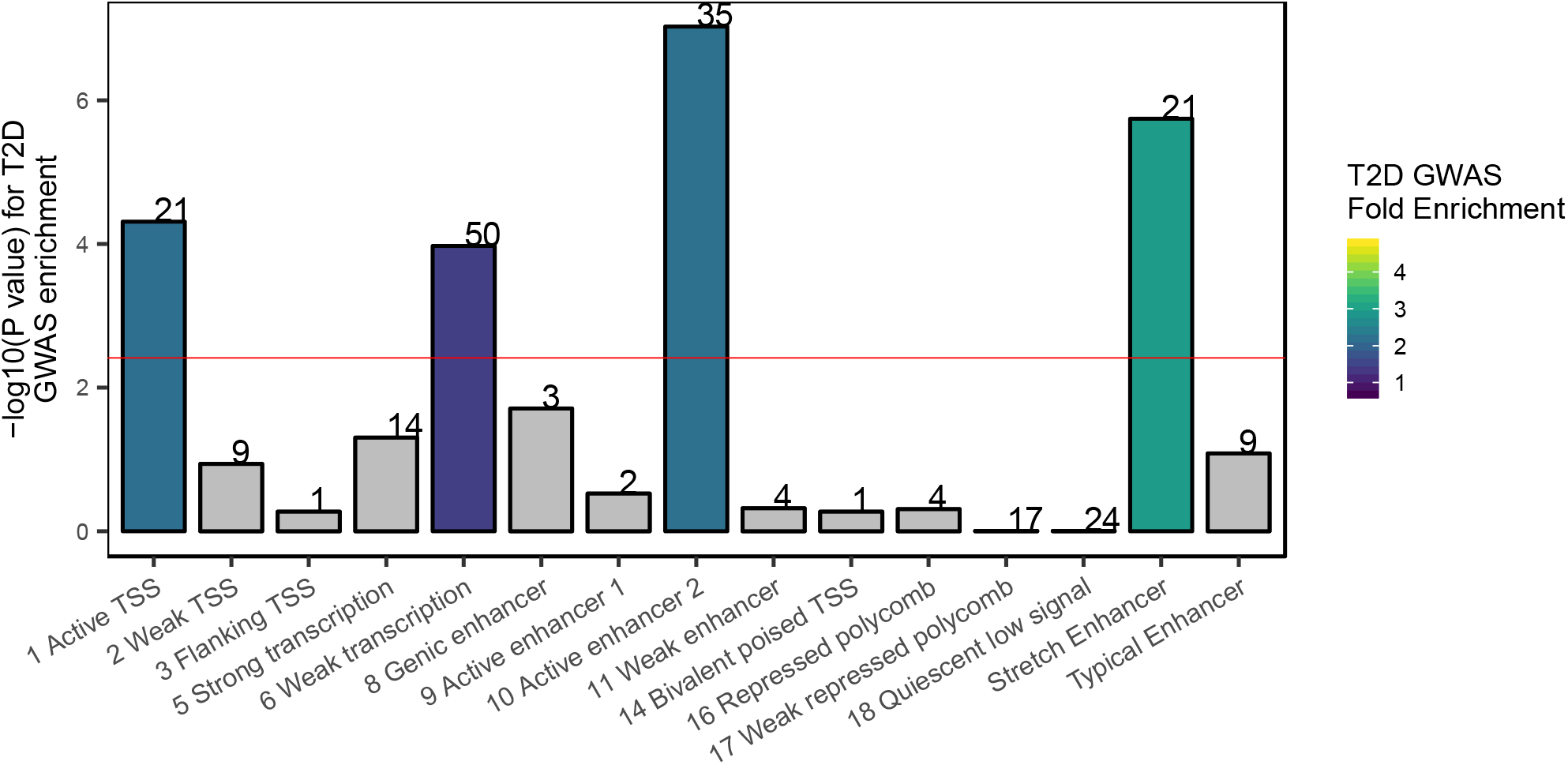
Fold Enrichment and –log10(P values) for T2D GWAS loci to overlap 13 chromatin states and stretch and typical enhancers. Labels indicate the number of independent T2D GWAS signals overlapping with each chromatin state. Red line = P value threshold after Bonferroni correction, adjusting for 15 tests. Grey = Not significant after Bonferroni correction

**Supplemental Figure SF7.**
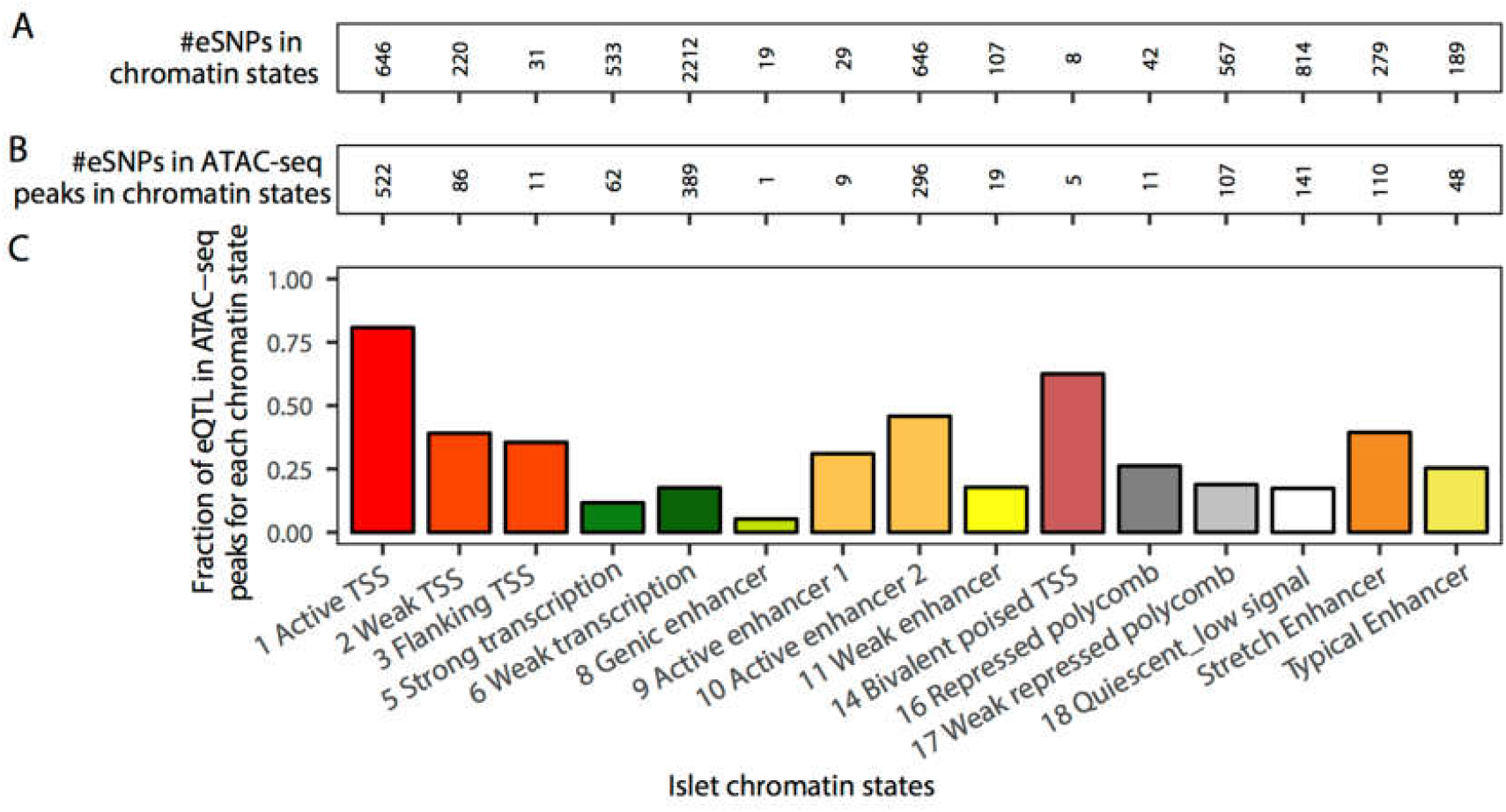
Fraction of eQTLs in ATAC-seq peaks in chromatin states. A: Number of eQTL Islet eQTL overlapping with Islet chromatin states and stretch/typical enhancers. B Number of Islet eQTL in Islet ATAC-seq peaks in chromatin states. C: Fraction of Islet eQTL in ATAC-seq peaks in each chromatin states. An eQTL overlap is considered if the eQTL lead eSNP or proxy SNP (LD r2>0.99) overlaps the feature.

**Supplementary Figure SF8.**
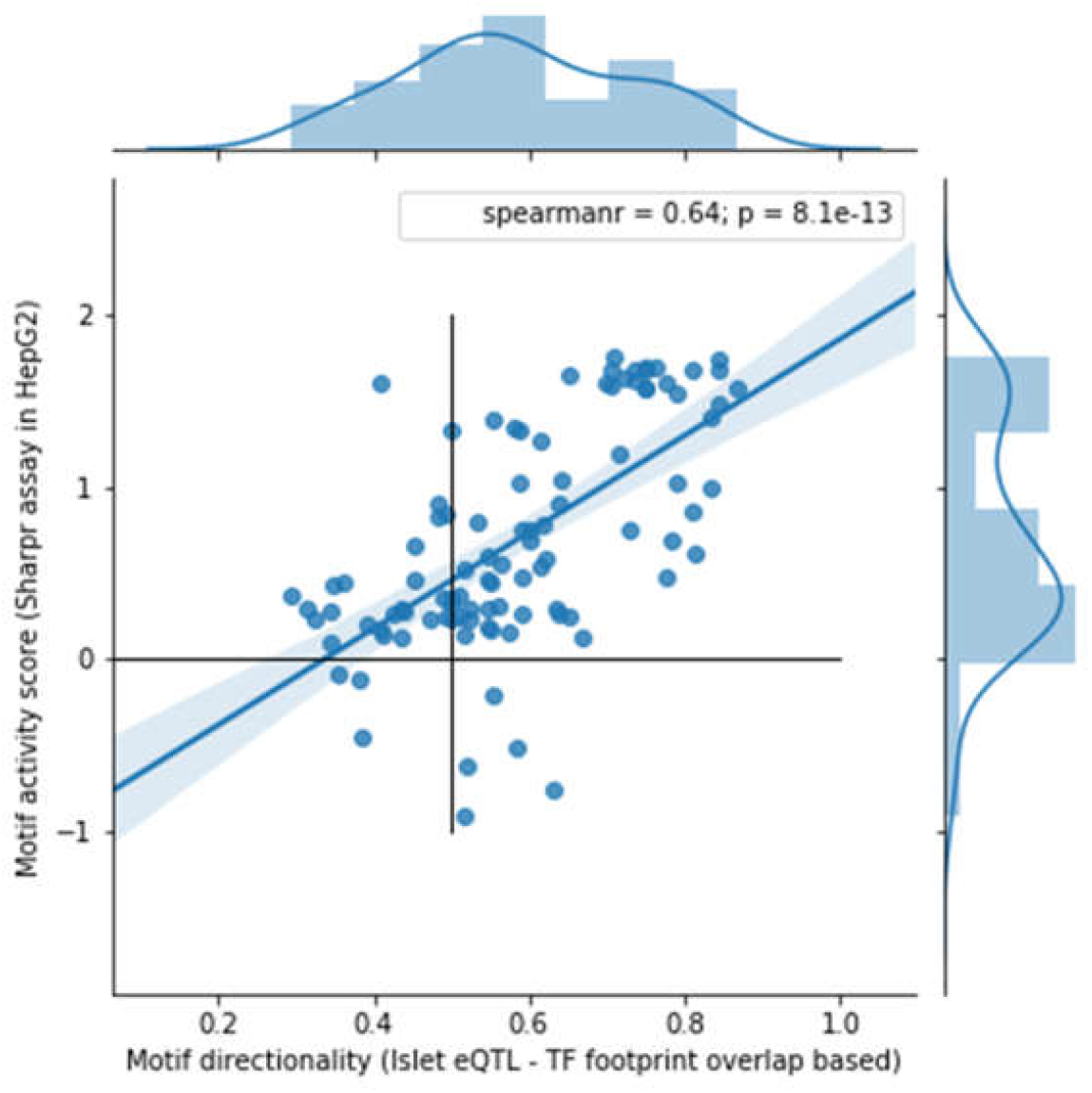
Comparison of MPRA. Transcription factor motif activity scores from Sharpr MPRA in HepG2 cells (CITE Kellis DOI 10.1038/nbt.3678) vs Motif directionality fractions from Islet eQTL and ATAC-seq TF footprinting data. TF Motifs that were reported to be either activating or repressive (P<0.01) from the MPRAs in both HepG2 and K562 are shown.

**Supplementary Figure SF9.**
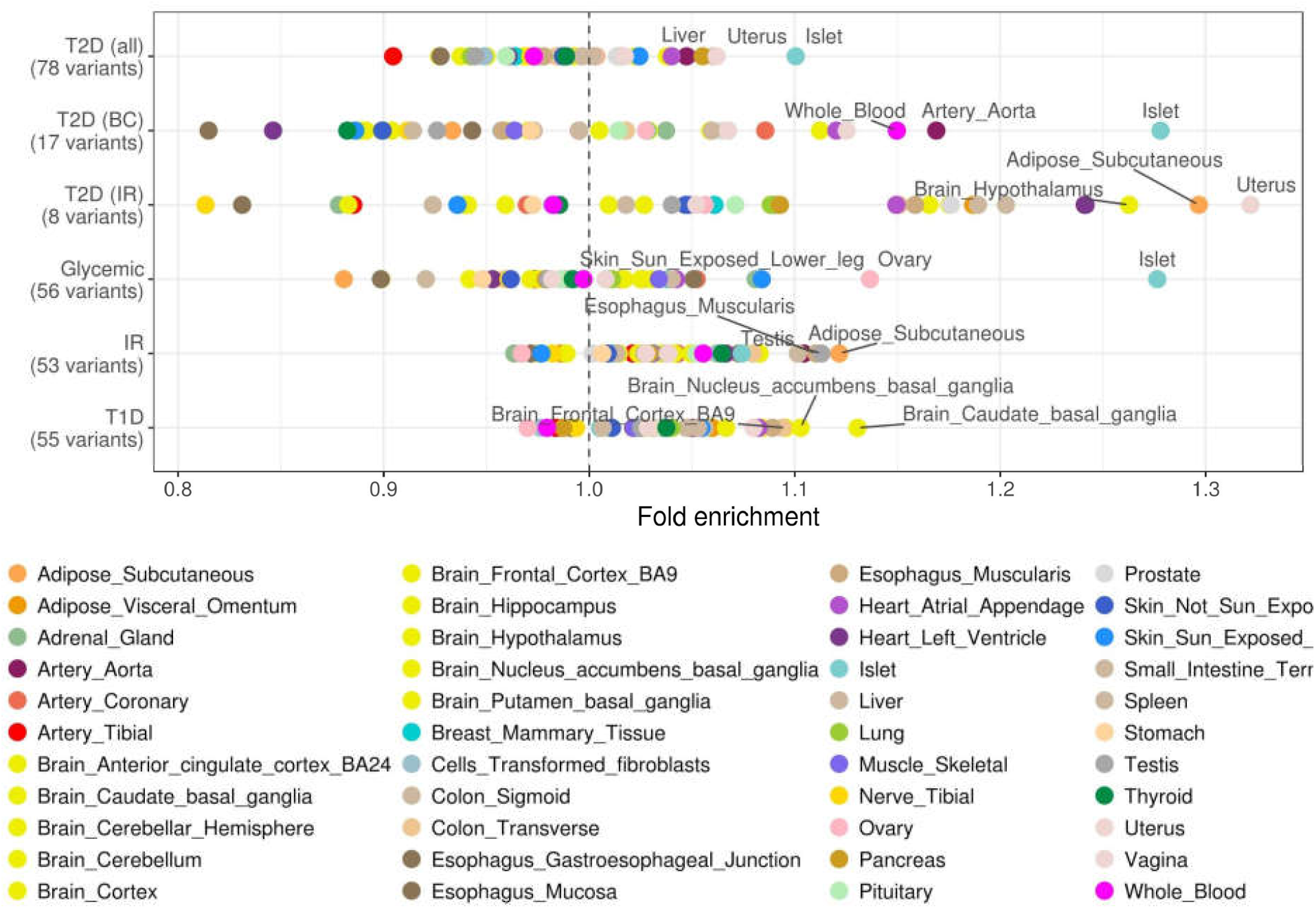
Tissue enrichment analysis in all GTEx tissues. Results support the conclusion show in the main figure that islets outperform other tissues for GWAS loci enrichment.

**Supplementary Figure SF10.**
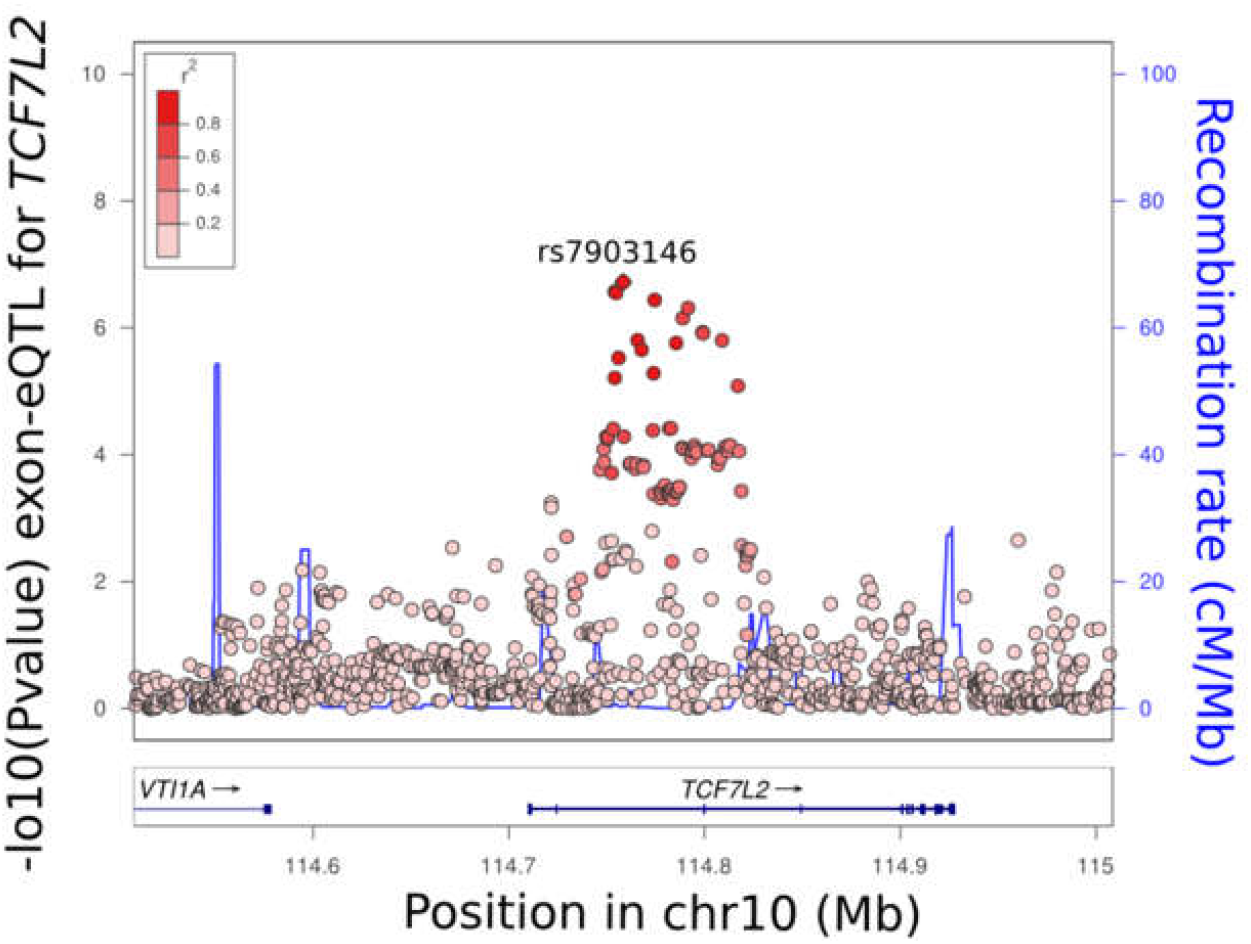
Locus-zoom showing the eQTL for *TCF7L2*.

**Supplemental Figure SF11.**
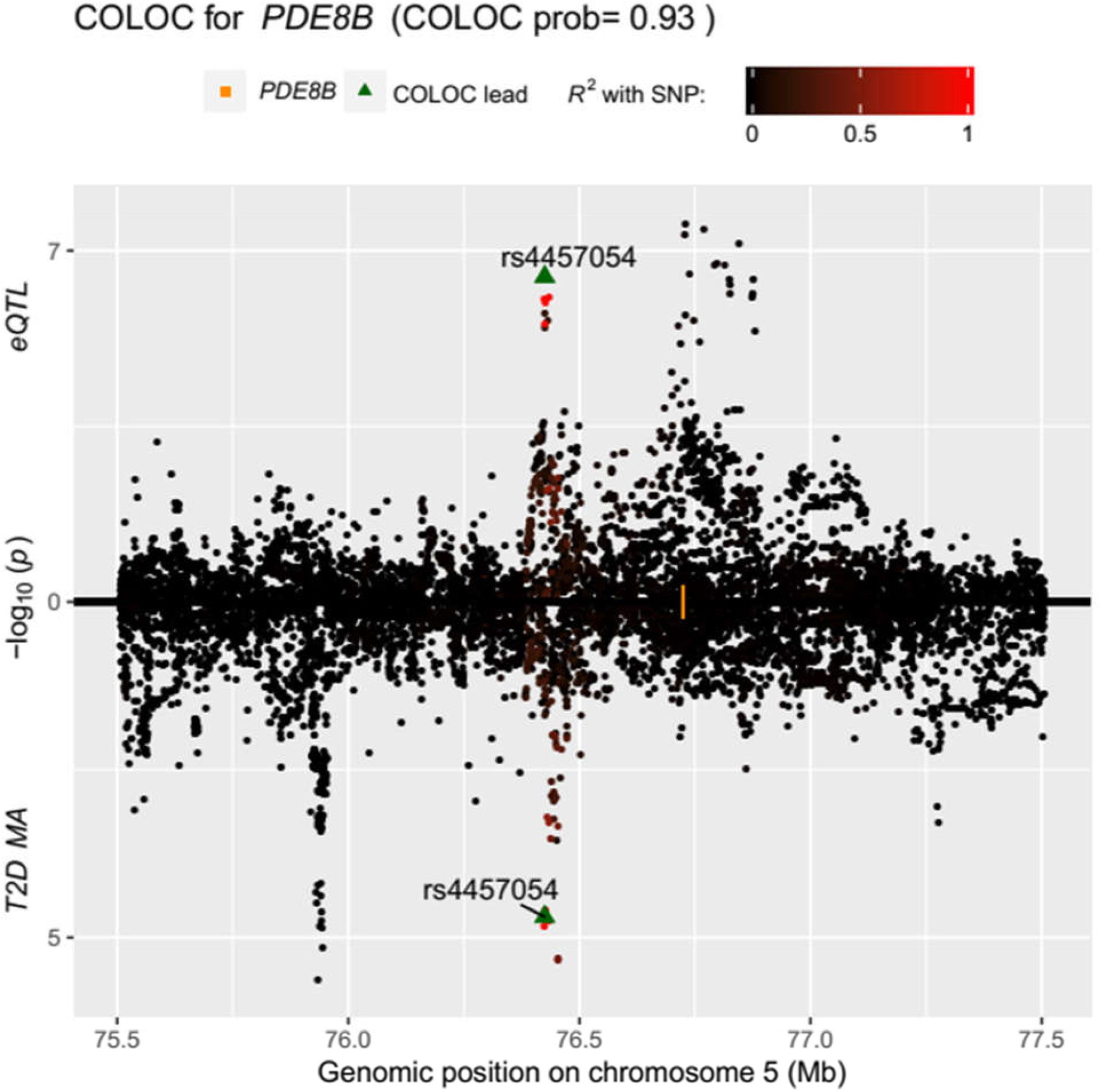
Miami plot of the eQTL for *PDE8B* on the top, and the GWAS for T2D on the bottom.

**Supplementary Figure SF12.**
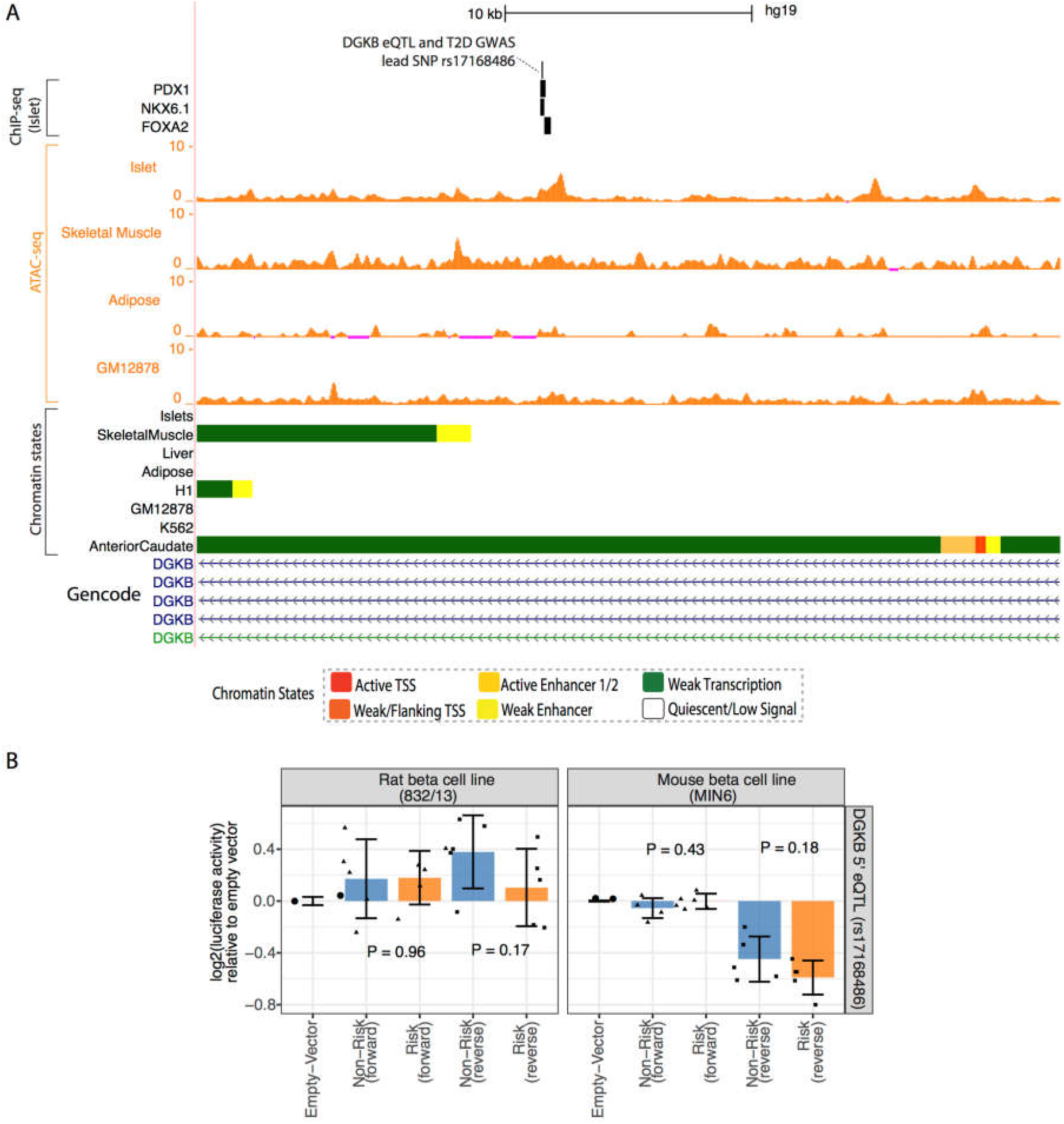
5’ eQTL (DGKB eQTL and T2D GWAS lead SNP 17168486). A: Genome browser shot of the 5’ DGKB eQTL along with ChIP-seq, ATAC-seq and chromatin state profiles in Islets and other tissues. B. Luciferase assay activities (normalized to empty vector) in rat (832/13) and mouse (MIN6) cell lines for the element containing the T2D GWAS and islet eQTL lead SNP (rs17168486), cloned in both forward and reverse orientation with respect to the *DGKB* gene. Differences between activities of the risk and non-risk allele containing elements were non-significant.

**Supplementary Figure SF13.**
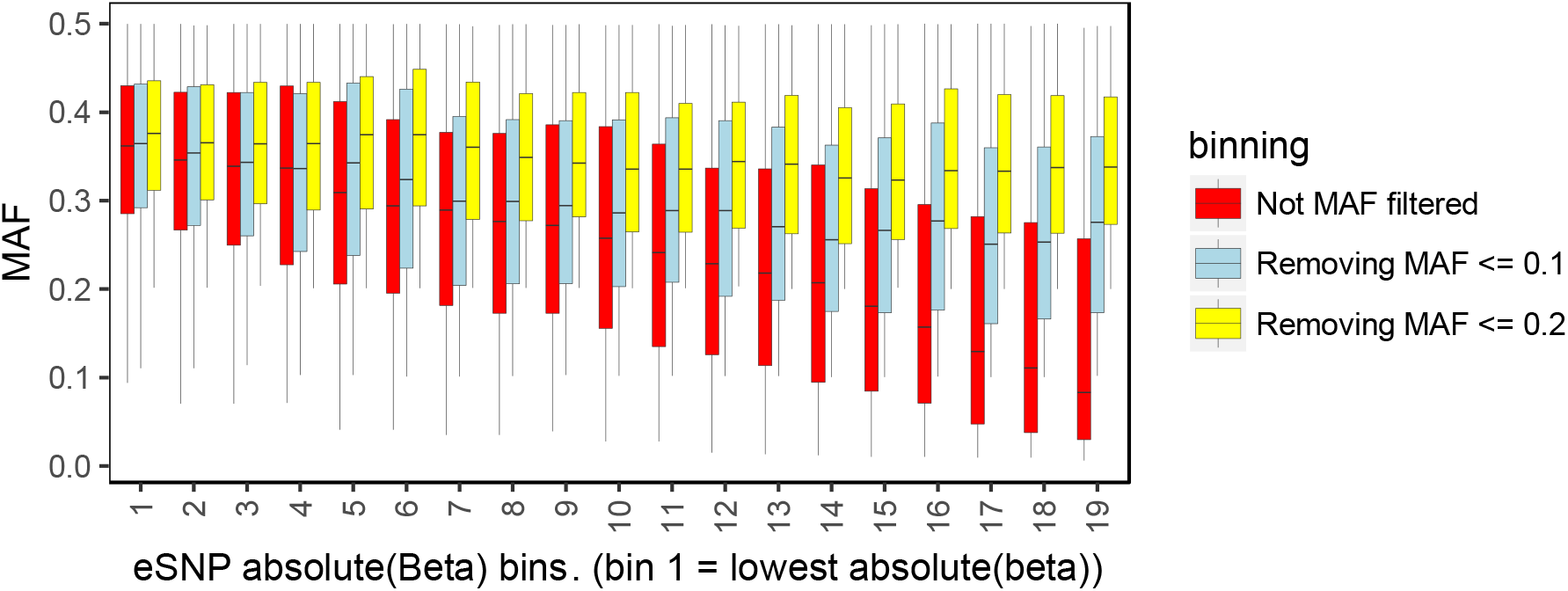
MAF filtering for eSNPs. MAF for islet eQTL eSNPs binned by absolute(beta) into equal sized, 50% overlapping bins. Bin 1 contains eSNPs with lowest absolute(beta), bin19 contains eSNPs with highest absolute(beta).

**Supplemental Figure SF14.**
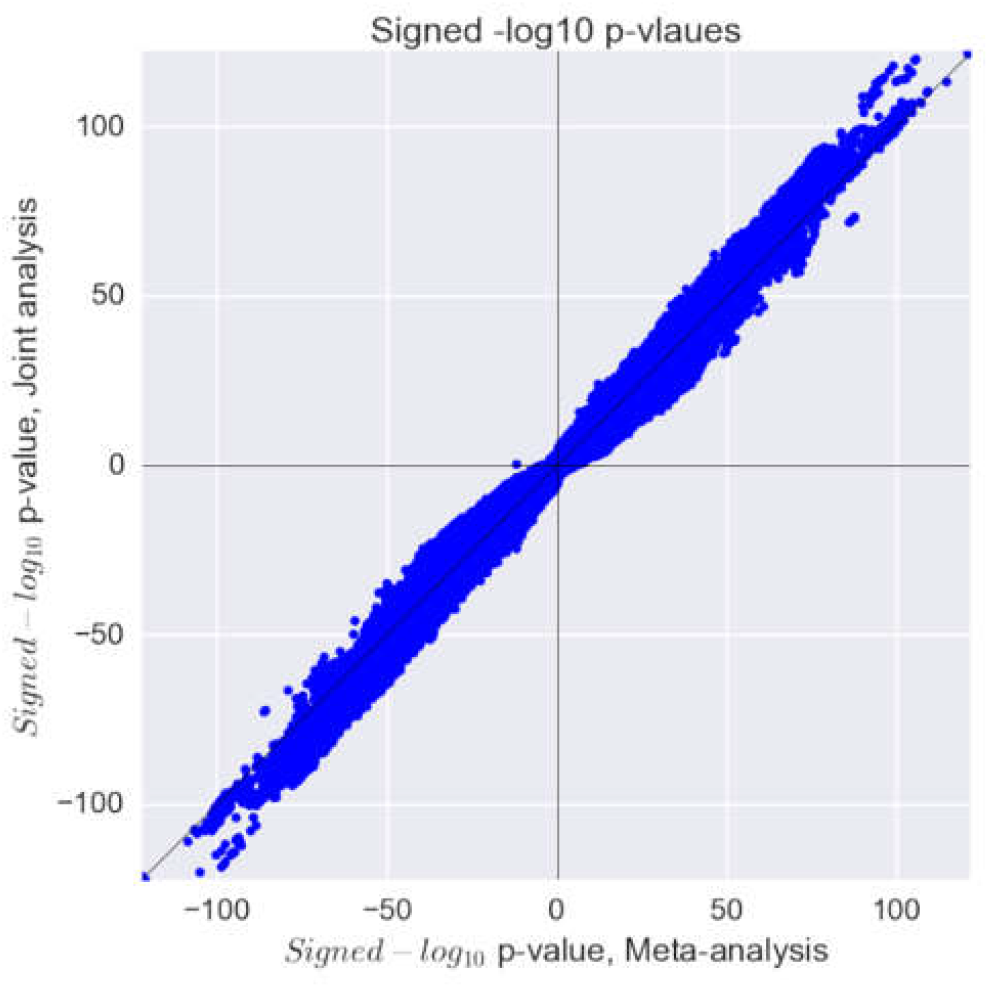
Comparison or meta-analysis of the four studies versus join re-processing and analysis. A comparison between our joint analyses and a fixed effects meta-analysis of the four studies found highly correlated results indicating appropriate control of the differences across studies

**Supplementary Figure SF15.**
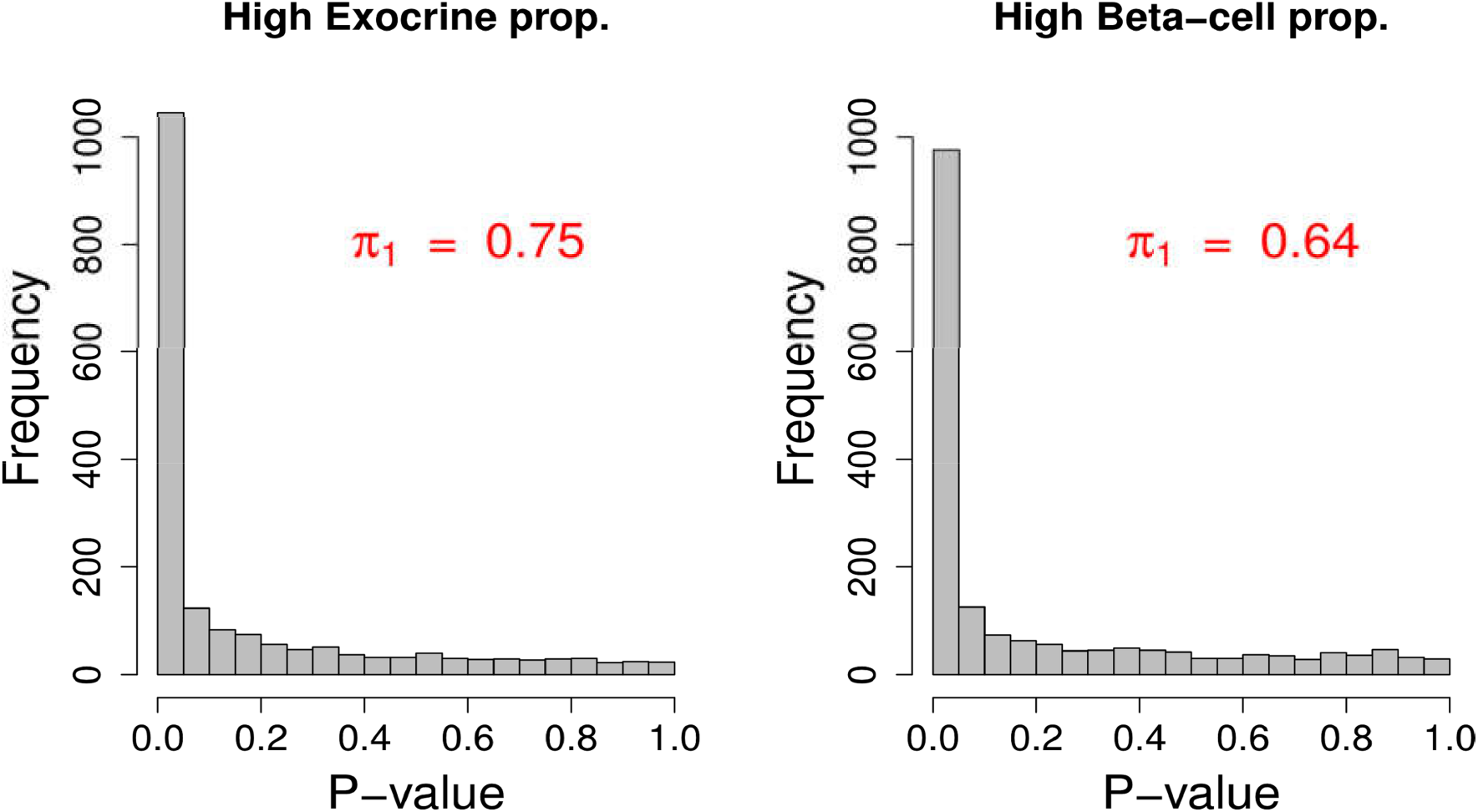
Replication rate of pancreas eQTLs in 100 islets with high proportion of exocrine expression (left) and in 100 islets with high proportion of beta-cells expression.

## Supplemental Tables list

ST1: List of lead eQTLs per gene in islets (exon level quantifications).

ST2: List of lead eQTLs per gene in islets (gene level quantifications).

ST3: List of 337 eQTLs only active in islets.

ST4: List of lead eQTLs per gene in beta-cells (exon level quantifications).

ST5: List of lead eQTLs per gene in GTEx pancreas (exon level quantifications).

ST6: List of 227 significant eQTLs in islets, also active in Beta-cells.

ST7: List of lead GxBeta-cells proportions eQTLs.

ST8: List of lead GxNonBeta-cells proportions eQTLs.

ST9: List of lead GxExocrine-cells proportions eQTLs.

ST10: List of bulk enrichment of footprints

ST11: Binned enrichment of footprints.

ST12: Motif directionality results.

ST13: List of GWAS variants used GWAS enrichment analysis.

ST14: Enrichment analysis results for GWAS-eSNPs across GTEx tissues.

ST15: Joint results for COLOC analysis with ENGAGE GWAS results for fasting glucose (FG) and DIAGRAM GWAS results for T2D.

ST16: Overlap between eQTL signals and glycemic response.

ST17: Reporter assay primers: Oligonucleotide primers used for the transcriptional reporter assays

ST18: EMSA probes: Oligonucleotide probes used for the electrophoretic mobility shift assays

