## Supplemental table 15 for "Influence of genetic variants on gene expression in human pancreatic islets – implications for type 2 diabetes"

| **Trait (Study)** | **Locus_Name** | **Locus_Index** | **Chr** | **Gene_Name** | **Strand** | **eSNP** | **R^2** | **Source** |
| --- | --- | --- | --- | --- | --- | --- | --- | --- |
| T2D | RBMS1 | rs7593730 | 2 | ITGB6 | - | rs764729 | 0.47 | COLOC+RTC |
| FG | SIX3-SIX2 | rs895636 | 2 | SIX3 | + | rs12712928 | 0.7 | RTC |
|  |  |  |  | SIX2 | - |  | 0.72 | RTC |
|  |  |  |  | RP11-89K21.1 | - |  | 0.7 | RTC |
|  |  |  |  | AC012354.6 | + |  | 0.72 | RTC |
| T2D; FG | ADCY5 | rs11708067 | 3 | ADCY5 | - | rs11708067 | 1 | COLOC+RTC |
| T2D; FG | IGF2BP2 | rs6769511 | 3 | IGF2BP2 | - | rs9854769 | 0.98 | COLOC+RTC |
| T2D | ST6GAL1 | rs16861329 | 3 | RPL39L | - | rs3887925 | 0.18 | RTC |
| T2D | UBE2E2 | rs1496653 | 3 | UBE2E2 | + | rs13094957 | 1 | COLOC+RTC |
| T2D; FG | ZBED3 | rs7708285 | 5 | PDE8B | - | rs335628 | 0.01 | COLOC |
| T2D; FG | DGKB (1) | rs6960043 | 7 | DGKB | - | rs10231021 | 0.94 | COLOC+RTC |
| T2D | DGKB (2) | rs17168486 | 7 | DGKB | - | rs17168486 | 1 | COLOC+RTC |
| FG | GRB10 | rs6943153 | 7 | GRB10 | - | rs6948959 | 0.29 | COLOC |
| T2D | GPSM1 | rs11787792 | 9 | CARD9 | - | rs61386106 | 0.69 | RTC |
| FG | IKBKAP | rs16913693 | 9 | CTNNAL1 | - | rs28361096 | 0.84 | COLOC+RTC |
| FG | ADRA2A | rs10885122 | 10 | ADRA2A | + | rs36106955 | 0.43 | RTC |
| T2D | CDC123-CAMK1D | rs11257655 | 10 | CAMK1D | + | rs11257658 | 0.94 | COLOC+RTC |
| T2D; FG | TCF7L2 | rs7903146 | 10 | TCF7L2 | + | rs7903146 | 1 | COLOC+RTC |
| T2D; FG | CENTD2-ARAP1 | rs11603334 | 11 | STARD10 | - | rs11603334 | 1 | COLOC+RTC |
| FG | FADS1 | rs174550 | 11 | FADS1 | - | rs61896141 | 0.32 | COLOC |
| FG | MADD | rs10501320 | 11 | MADD | + | rs11039165 | 0.95 | COLOC+RTC |
| T2D | KLHDC5 | rs10842994 | 12 | KLHL42 | + | rs4931479 | 0.03 | COLOC |
| T2D | MPHOSPH9 | rs1727313 | 12 | CDK2AP1 | - | rs6576 | 0.75 | RTC |
| T2D | ANK1 | rs516946 | 14 | NKX6-3 | - | rs4736819 | 0.27 | RTC |
| FG |  | rs12549902 |  |  | - |  | 0.93 | RTC |
| FG | WARS | rs3783347 | 14 | WARS | - | rs2146105 | 0.95 | RTC |
| T2D | AP3S2 | rs2028299 | 15 | AP3S2; C15orf38-AP3S2 | - | rs2165069 | 0.96 | RTC |
| T2D | HMG20A | rs7177055 | 15 | HMG20A | + | rs59153558 | 0.95 | COLOC+RTC |
