## Supplemental table 17 for "Influence of genetic variants on gene expression in human pancreatic islets – implications for type 2 diabetes"

Table: Oligonucleotide primers used for the transcriptional reporter assays.

| **Primer** | **Sequence (5’-3’)** | **Amplified region (hg19)** |
| --- | --- | --- |
| Haplotype A_Forward  Haplotype A_Reverse | GAGTTGCTGGAATGGAGGAG  CATGCGTCGCAATGTCTTT | chr7:15,055,310-15,056,074 |
| Haplotype B_Forward  Haplotype B_Reverse | TGCTAAGGCTAACCATAAACCA  TCAGCCAGCTGAAAGCTAGA | chr7:15,064,030-15,064,621 |
