## Supplemental table 18 for "Influence of genetic variants on gene expression in human pancreatic islets – implications for type 2 diabetes"

Table: Oligonucleotide probes used for the electrophoretic mobility shift assays.

| **Variant** | **Sequence (5’-3’) ‡** | **Position** |
| --- | --- | --- |
| rs10228796 | ACTGGGTT(**C/G)**TACCTTAA | chr7:15064182-15064198 |
| rs10258074 | TGGCAACC**(A/T)**GTCTCCTA | chr7:15064208-15064224 |
| rs2191348 | TTACCAAT**(G/T)**CAATACCA | chr7:15064247-15064263 |
| rs2191349 | AGGATCAT(**G/T)**TAAGACCA | chr7:15064301-15064317 |

**‡**Variant alleles are in bold.
